# Diversification of molecularly defined myenteric neuron classes revealed by single cell RNA-sequencing

**DOI:** 10.1101/2020.03.02.955757

**Authors:** Khomgrit Morarach, Anastassia Mikhailova, Viktoria Knoflach, Fatima Memic, Rakesh Kumar, Wei Li, Patrik Ernfors, Ulrika Marklund

## Abstract

Autonomous functions of the gastrointestinal tract require the combined activity of functionally distinct neurons of the enteric nervous system (ENS). However, the range of enteric neuron diversity and how it emerges during development remain largely unknown. We here make a novel molecular definition of 12 enteric neuron classes (ENCs) within the myenteric plexus of the mouse small intestine. We identify communication features and provide histochemical markers for discrete motor, sensory, and interneurons together with genetic tools for class-specific targeting. Transcriptome analysis of embryonic ENS reveals a largely post-mitotic principle of diversification, where only ENC1 or ENC8 phenotypic traits arise through a binary neurogenic trajectory, and other identities form through subsequent differentiation. We propose generic and class-specific transcriptional regulators and functionally connect the transcription factor *Pbx3* to one post-mitotic identity conversion. Our results offers a conceptual and molecular resource for dissecting ENS circuits, and predicting key regulators for the directed differentiation of distinct enteric neuron classes.

## INTRODUCTION

Within the gastrointestinal (GI) tract, the enteric nervous system (ENS) controls motility, blood flow and secretion, but also communicates with the immune system and microbiome.^1,2^ These physiological processes rely on the combined activity of diverse enteric neuron types. Meticulous work over the last decades has identified a framework of enteric neurons with sensory (Intrinsic Primary Afferent Neurons, IPANs), motor or interneuron characteristics.^3^ Notably, only a few studies have focused on the mouse, despite its primary role as experimental animal.^4,5,6^ To make ENS attainable to modern genetic technologies, the unique molecular signatures of the full range of mouse enteric neurons would be desirable. For instance, this could resolve whether the multiple roles of IPANs (first neurons in intrinsic reflex circuits) rely on subtypes with different properties, information that would help therapy development for gut motility disorders targeting this neuron type.^7^

That functional diversity amongst enteric neurons is critical for physiological homeostasis is evidenced in several gut disorders where selective neuron types are affected, including esophageal achalasia (reduced NOS1^+^ neurons).^8,9^ Likewise, the absence of all enteric neurons in the distal bowel of Hirschsprung disease patients causes chronic constipation and requires surgical resection. Motivated by the lack of satisfactory treatments for enteric neuropathies, and spearheaded by the recent derivation of enteric neurons from human pluripotent stem cells (PSC)^10^, efforts are now placed on developing reparative cell therapies.^11^ Potential approaches to repopulate ENS-deficient bowel segments include transplantation of *in vitro*-derived stem cells and steering of endogenous stem cells to neurons. In either case, complete functional recovery will depend on the success of recreating a correct cellular ENS composition. This emphasizes the importance of acquiring a refined picture of the neuron subtypes that make up the normal ENS, and how they specialize during development.

The ENS is mainly generated from vagal neural crest cells that colonize the foregut at embryonic day (E)9 and reach the hindgut at E14 where they converge with sacral-derived neural crest.^12^ Subsequently, Schwann cell precursors (SCPs), associated with extrinsic nerves, invade the gut wall and contribute to the ENS.^13^ Unlike the developing central nervous system (CNS), where neuron subtypes emerge at stereotypical positions from patterned zones of stem cells^14^, the ENS develops from highly motile neural crest-derived streams of intermingled enteric stem cells and differentiating glia and neurons.^12^ Phenotypically different neurons undergo neurogenesis at different embryonic time windows, suggesting a temporal mode of diversification.^15,16^ The temporo-spatial expression of transcription and signaling regulators in the developing ENS has been comprehensively mapped, but this atlas lacked refinement to describe lineage development of distinct neuron classes.^17^

In this study we optimized efficient capture of enteric neurons for single cell transcriptomics and evidence a novel taxonomy of 12 myenteric neuron classes of the mouse small intestine, including motor, inter- and sensory neuron types, largely expanding and refining our earlier characterisation.^18^ The Enteric Neuron Classes (ENCs) are defined by their communication (signaling, synaptic and connectivity) features, which we portray along with immunohistochemical markers and genetic tools for immediate use in experimental neuroscience. Based on further transcriptome analysis of the developing ENS we postulate a new logic for ENS diversification where myenteric ENC identities are generated from an initial binary difference further diversified at the post-mitotic state. The principle is supported by the finding of a transcription factor (*Pbx3*) that regulate the post-mitotic conversion of immature NOS1^+^/GAL^+^/VIP^+^ ENC8/9 neurons into CALB^+^ ENC12 neurons.

## RESULTS

### Single cell RNA-sequencing reveals 12 Enteric Neuron Classes in the myenteric plexus of the mouse small intestine

We have previously reported single cell RNA sequencing (scRNA-seq) of myenteric plexus isolated from the small intestine of *Wnt1-Cre;R26R-Tomato* mice, yielding 1105 neurons classified into nine clusters. We suspect that the number of neurons was insufficient to represent the full range neuronal diversity of the ENS. Seeking to enrich neuron proportion, we investigated whether the recently described pan-neuronal *Baf53b-Cre* mice^19^ specifically and efficiently labels enteric neurons, when mated to *R26R-Tomato* reporter mice. We quantified co-expression of TOM with the neuron marker HUC/D and the enteric glia markers SOX2/SOX10 in myenteric plexus across different regions of the small intestine. We found that all TOM^+^ cells expressed HUC/D whereas none expressed SOX2/10 (Fig. 1a). Likewise, the vast majority of HUC/D^+^ cells expressed TOM (on average 98%; Fig 1b). An improved protocol (see material and methods) for cell dissociation was applied on the myenteric plexus from the small intestine of *Baf53b-Cre;R26R-Tomato* mice, TOM^+^ cells were sorted by flow cytometry and single cells analyzed by 10 x Chromium RNA sequencing (scRNA-seq)(Fig. 1c).

**Figure 1.**
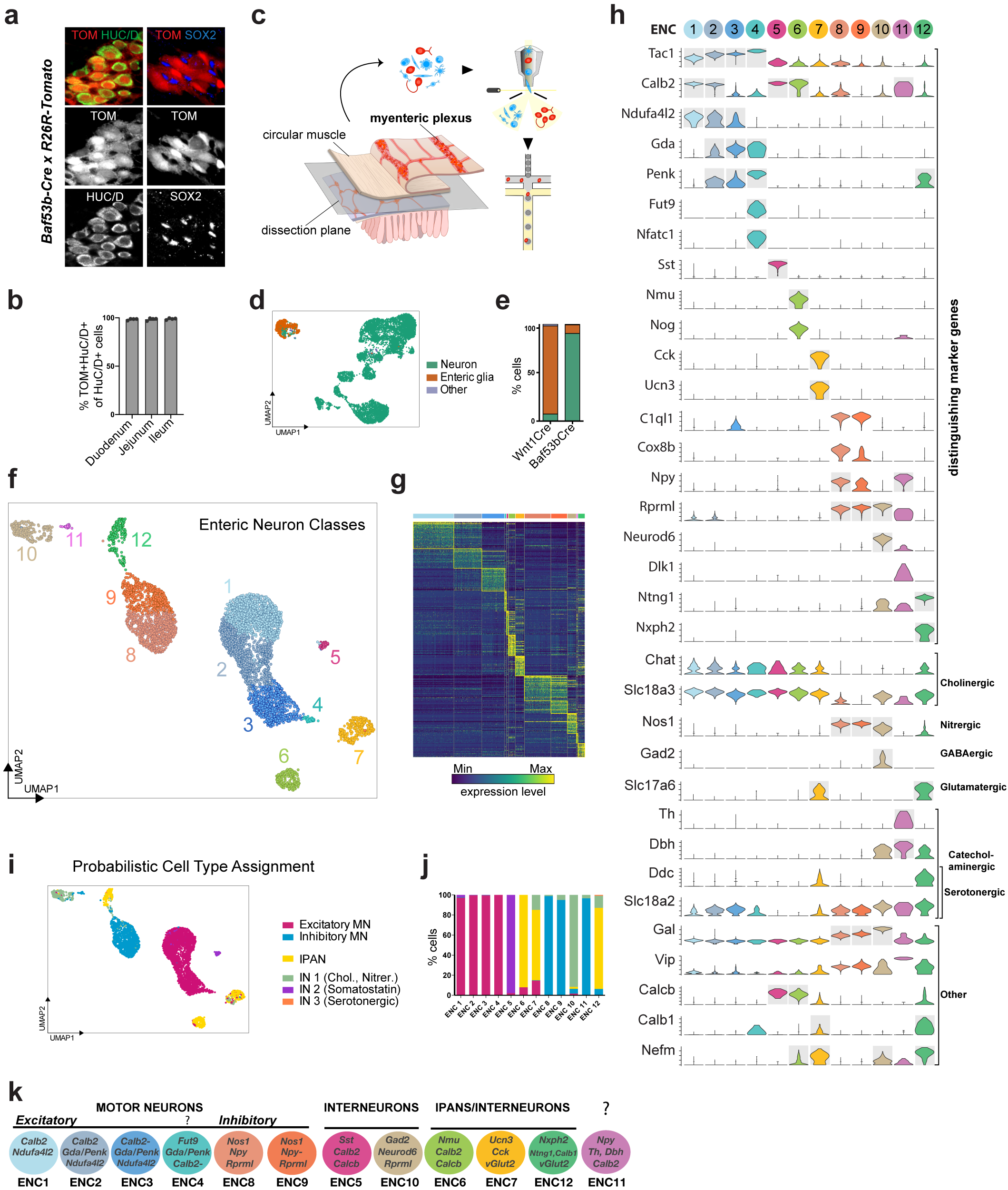
Molecular definition of 12 Enteric Neuron Classes (ENCs) in the mouse small intestine myenteric plexus. **a)** Peels of the myenteric plexus from *Baf53b-Cre;R26R-Tomato* mice at postnatal day (P)26, showing Tomato (TOM, red) expression in enteric neurons (HUC/D^+^, green) but not in enteric glia (SOX2^+^, blue). **b)** Graph showing the average percentage of HUC/D^+^ cells expressing TOM within the duodenum (2138 cells), jejunum (2791 cells) and ileum (3180 cells); n=4 at P24-26. TOM was not detected in any SOX2^+^/SOX10^+^ cells (1848 cells counted in 3 animals; data not in graph). **c)** Schematic drawing of experimental procedure indicting the dissection plane to retrieve myenteric plexus of *Baf53b-Cre; R26R-Tomato* small intestine, flow sorting of single TOM^+^ cells followed by 10x chromium single cell RNA-sequencing. **d)** UMAP representation of sequenced cells with total UMI counts > 600, annotated by probabilistic cell type assignment (neuron, glia, other) based on priori markers. **e)** Comparison of the proportion of major cell types in current single cell dataset obtained from *Baf53b-Cre;R26R-Tomato* mice and our published dataset obtained from *Wnt1-Cre;R26R-Tomato* (Zeisel et al, 2018). **f)** Unsupervised clustering of 4892 high quality enteric neurons, color-coded by enteric neuron classes (ENC)1-12 and represented on UMAP. **g)** Heatmap representing top 50 (by average logFC) differentially expressed genes of each ENC. **h)** Violin plots representing the expression (log-scale) of key genes among ENCs, including new class-specific marker genes and previously discovered neurotransmitter/neuropeptide genes. Grey boxes indicate genes whose expression were confirmed by immunohistochemistry (see Fig. 3); together they form combinatorial codes that are unique for each ENC. A gene set including *Calb2* and stronger *Ndufa4l2* demarcated ENC1-2 from ENC3-4, while *Gda* and *Penk* expression discriminated ENC2-4 from ENC1. ENC4 resembled ENC3, but displayed a unique expression of *Fucosyltransferase 9* (*Fut9*) and the transcription factor *Nfatc1*. Expression of selected markers genes are also represented on UMAPs (see Extended Data Fig. 2). **i)** Probabilistic neuron subtype assignment based on literacy-curated functional type markers (see Material and Methods) represented on UMAP. **j)** Proportion of learned functional neuron subtypes in each ENC. **k)** Schematic drawing indicating proposed functional assignment and selected combinatorial marker genes for each ENC. ENC: Enteric Neuron Class; FC: Fold Change; MN: Motor neuron; IN: Interneuron; UMAP: Uniform Manifold Approximation and Projection; UMI: Unique Molecular Identifiers;

Of 9,141 cells captured we retained those with 600 total unique molecular identifier (UMI) counts to assess the proportion of neurons in the dataset (Fig. 1d). More than 90% of the recovered cells corresponded to enteric neurons, compared to only 10% in our previous capture from *Wnt1-Cre;R26R-Tomato* mice^18^ (Fig. 1e). We employed iterative clustering with increasingly restrictive quality control thresholding, and removed low-quality, non-neuronal and ambiguous clusters. We retained 4,892 high quality enteric neurons for unsupervised graph-based clustering, yielding 12 enteric neuron classes (ENCs) (Fig. 1f,g; Extended Data Fig. 1a). Note that our new classification confirmed 4 of the previously found clusters, re-defined 6 clusters and identified 2 new distinct clusters (Extended Data Fig. 1b).

We discovered the most highly enriched genes in each class (Fig. 1g,h). Notably, some classes were defined by the selective expression of single genes with likely importance for their neuronal activity. These included ENC5 (*somatostatin*; *Sst*), ENC6 (*neuromedin U*; *Nmu*), ENC7 (*cholecystokinin*; *Cck* and *urocortin 3*; *Ucn3*), ENC10 (*Neurod6*) and ENC12 (*neurexophilin-2*; *Nxph2* and enriched *netrin-G1*; *Ntng1*). Other classes were distinguished by the combinatorial codes of enriched gene expression (Fig. 1h; see Feature Plots in Extended Data Fig. 2a,b). ENC1-4 displayed a partly shared gene expression, including *Tac1*, but differed in their expression of *Calb2, Ndufa4l2, Gda, Penk* and *Fut9* (Fig. 1h). ENC8-9 jointly expressed *Nos1* and *c1ql1*, but *Npy* and *Cox8b* were significantly lower in ENC9. ENC11 lacked *Nos1* but displayed highest *Npy* expression of all ENCs.

We next sought to relate our new ENS classification to previously discovered characteristics of functional enteric neuron types. To this end, we gathered binary phenotypic marker codes from a series of immunohistochemical studies of mouse myenteric neurons,^4,5,6^ performed probabilistic cell type assignment using these markers as priori information (Fig. 1i) and regrouped according to the unsupervised ENCs. The ENCs generally consisted of cells with the same functional type (Fig. 1i,j). ENC1-4 were assigned to excitatory motor neurons (*Tac1/Calb2*), while ENC8 and 9 matched inhibitory motor neurons (*Nos1/Gal/Vip/Npy)*. ENC6, 7 and 12 were mapped to IPANs and expressed different combinations of previously used IPAN markers (*Calca/Calcb/Nfem/Calb1/Calb2)*. These plausible IPANs were clearly separable by their selective expression of neuroactive substances *Nmu*, *Ucn-3/Cck* or *Nxph2*. The IPAN identity of ENC6 and 12 was confirmed, while ENC7 likely have different functions (See Fig. 4). ENC10 was assigned to Interneuron (IN)1, and was, contrary to most enteric neurons, both nitrergic (*Nos1*) and cholinergic (*Slc18a3; Chat* detection level overall low^18^). ENC5 expressed *Sst/Calcb/Calb2* and was mapped to IN2. Characteristics of serotonergic IN3, (*Ddc, Slc18a2, Slc6a4*, Fig. 1h; Extended Data Fig. 2c) were detected in a subset of ENC12 cells.

Gene profiles signified several additional neuroactive substances with class-specific expression. *vGlut2* and *Gad2* expression suggested glutamatergic (ENC7,12) and GABAergic (ENC10) phenotypes (Fig. 1h). ENC11 expressed *Dbh* and *Th,* key enzymes in the noradrenaline biosynthesis (Fig. 1h), and could therefore represent a new neuron type (Fig. 1k).

### Enteric Neuron Classes are defined by their unique communication features

Synaptic input-output communication represents a fundamental distinction among neuron types and is determined by connectivity, responsiveness, and release of neurotransmitters/peptides.^20^ We explored gene families that control these functions and found differential expression between the ENCs (Extended Data Fig. 3, Fig. 2a-d). Responsiveness to enteric neurotransmitters/peptides appeared class-specific. For example, glutamate receptor *Grm5* was selectively expressed in ENC7, while somatostatin and Npy receptors *Sstr5* and *Npy5r* were limited to ENC12 (Fig. 2a). Distinct expression of many cell signaling molecules with unknown functions in the mature ENS (e.g. *Edn1*, *Kit*, *Bdnf*) was also detected amongst ENCs. Notably, ENC6 selectively expressed signaling factors/modulators recently linked to enteric control of immune cells (*Nmu* and the BMP inhibitor *Nog*)^21–23^ (Fig. 1h, Extended Fig. 3b). Ion channels showed class-specific distribution likely reflecting unique electrophysiological properties (Fig. 2b; Extended Data Fig. 3e). ENC6 expressed *Kcnn3* and *Ano2*, described in sensory transduction^7,24^, while ENC12 expressed the mechanosensory channel *Piezo2*^25^, further supporting sensory identities of these two classes. ENCs also displayed unique combinations of adhesion molecules (e.g. semaphorins, cadherins and ephrins) that may determine their precise connectivity patterns (Fig. 2c; Extended Data Fig. 3d). Of notice, neurexophilins (*Nxph1-4*), modulators of synaptic plasticity^26^, displayed almost mutually exclusive expression patterns. Taken together, the selective expression of genes conferring neuronal properties strongly supports that ENC1-12 represent functionally distinct neurons.

**Figure 2.**
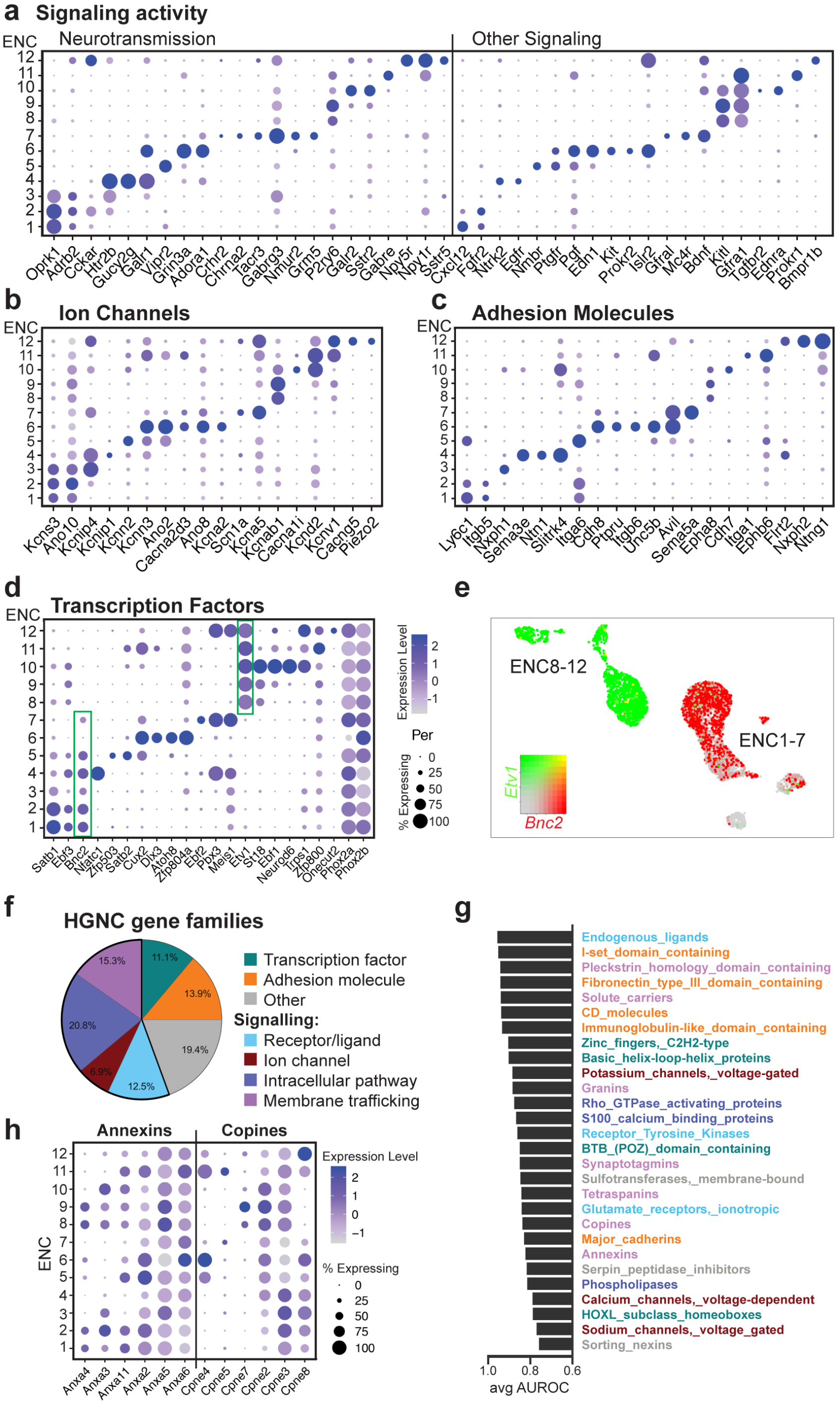
Gene categories conferring functional characteristics in Enteric Neuron Classes. **a-d)** Dot plots showing differentially expressed genes categorized as signaling molecules, ion channels, adhesion molecules and transcription factors. Color scale represents z-score and dot size represent percentage of cells with non-zero expression within a given class. See Extended Data Fig. 3 for dot plots of all enriched gene members within these gene categories. Green boxes in **(d)** highlight *Bnc2* and *Etv1* whose expression broadly span class 1-7 and 8-12, respectively. **e)** *Bnc2* and *Etv1* expression in individual cells depicted on UMAP. Color map represents scaled expression of *Bnc2* (red), *Etv1* (green), co-expressed (yellow) and non-expressing (grey). **f)** Pie chart showing the proportion of different categories of gene families with mean AUROC > 0,75. **g)** Selected top scoring HGNC gene families ordered by mean AUROC and color-coded according to gene family categories. See Supplementary Table 1 for the full list of the gene families. Top-scoring gene families corresponded to 69% of gene families defining diversity amongst cortical inhibitory neurons (Paul et al., 2017). **h)** Dot plot showing differentially expressed genes within two gene families categorized to membrane trafficking (see also Extended Data Fig. 3f). AUROC: area under the receiver operator characteristic curve; ENC: Enteric Neuron Class; HGNC: HUGO Gene Nomenclature Committee; UMAP: Uniform Manifold Approximation and Projection

Gene profiles conferring neuronal characters result from fine-tuned transcriptional programs. Interrogation of transcription factor expression revealed that each ENC could be distinguished by combinatorial sets of transcription factors (e.g. ENC6 (*Atoh8, Dlx3*), ENC7 (*Pbx3, Ebf2*), ENC10 (Neurod6), ENC12 (*Etv1, Pbx3, Onecut2*), (Fig. 2d; Extended Data Fig. 3c). ENC6 showed the most distinct profile and was the only class that lacked *Phox2a. Etv1* and *Bnc2* were expressed across several ENCs but with a complementary pattern relative to each other, possibly reflecting early developmental divisions (Fig. 2d,e).

To identify additional gene families that define ENC identities, we assessed HGNC (HUGO Gene Nomenclature Committee) gene-sets curated by Paul et al (2017)^20^ (Supplementary Table 1) in our ENCs, by calculating their mean AUROC (Area Under the Receiver Operating Characteristic curve) score. The majority of enriched gene families could be categorized according to their transcription, adhesion and signaling activities (Fig. 2f,g). We also categorized 15% of the gene-sets to have “membrane trafficking” properties, including the relatively unexplored tetraspanins^27^, copines^28^, annexins^29^ (Fig. 2h, Extended Data Fig. 3f). Thus, our data suggest that proteins involved in organizing membrane distribution of receptors/channels/vesicles contribute to neuron subtype specific features.

### Identification of immunohistochemical markers for detection of Enteric Neuron Classes

To validate the existence of enteric neuron classes *in vivo* we performed immunohistochemistry in the mouse small intestine and included both positive (Fig. 3) and negative markers (Extended Data Fig. 4) for all ENCs. A total of 23 proteins were analysed in respect to our newly defined ENCs. Several antibodies for the same antigen and ENC were assessed to ensure faithful histochemical reagents for individual ENCs (Supplementary Table 2). Of notice, proteins previously not linked to distinct enteric neuron subtypes in the mouse ENS were detected including NDUFA4L2 (ENC1-3), GDA (ENC2-4), FUT9 (ENC4), NMU (ENC6), CCK, UCN3 (ENC7), RPRML (ENC8-11), NEUROD6 (ENC10), and NXPH2 (ENC12). Importantly, their expression were found in the context of a combinatorial marker code (See Extended Data Fig. 4p for a summary table). For instance, NEUROD6 readily co-expressed NOS1 and NF-M (Fig. 3, ENC10) but never co-appeared with CALR (Extended Data Fig. 4l).

**Figure 3.**
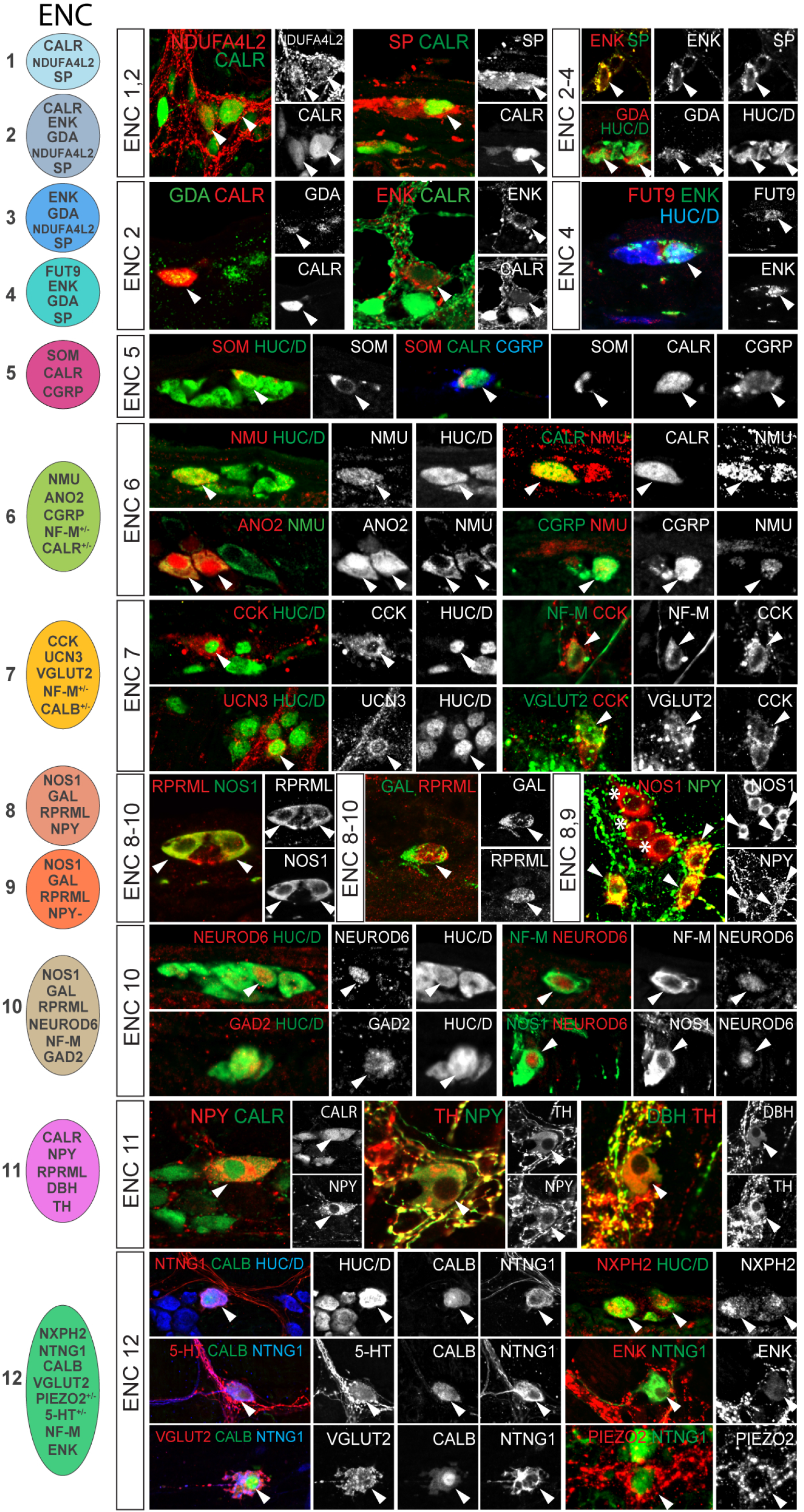
Immunohistochemical validation of marker genes for Enteric Neuron Classes. Immunohistochemical localization of selected key marker proteins of ENC1-12 in the small intestine myenteric plexus at P21-90. Pictures show either myenteric peel preparations or transverse sections. Images for CCK, VGLUT2 and CGRP were obtained from colchicine treated tissues where proteins are concentrated to neuron bodies. Arrowheads indicate co-expression of proteins. Stars indicate NOS1^+^ neurons lacking NPY expression (ENC9). Schematic cell drawings to the left indicate positive markers tested. See Extended Data Fig. 4 for negative marker genes and summary of all validated marker genes. CALB: Calbindin; CALR: Calretinin; CCK: Cholecystokinin; CGRP: Calcitonin Gene Related Peptide; DBH: Dopamine Beta Hydroxylase; ENK: Enkephalin; GAD2: Glutamic Acid Decarboxylase 2; GAL: Galanin; HUC/D: Elav-Like Protein 3/4; NF-M: Neurofilament M; NMU: Neuromedin U; NOS1; Nitric Oxide Synthase 1; NPY: Neuropeptide Y; NTNG1: Netrin G1; RPRML: Reprimo-Like; SOM: Somatostatin; SP: Substance P; TH: Tyrosine Hydroxylase; VGLUT2: Vesicular Glutamate Transporter 2

Analysis of protein expression enabled the additional study of neurotransmitters, which are catalytic aminoacid products and therefore not detectable in transcriptomes. Of notice, 5-HT (serotonin) was identified in a subset of ENC12 neurons (Fig. 3) as suggested from the cell assignment (Fig. 1i,j; Extended Data Fig. 2c). The analysis also supported GABAergic (GAD2), glutamatergic (VGLUT2) and noradrenergic (DBH, TH) phenotypes (Fig. 3, Fig 1k).

### ENC6 and subsets of ENC12 neurons show morphological and projectional IPAN traits

We choose to further investigate the three predicted IPANs (ENC 6,7 and 12) given the key functions neurons with sensory modulaties are expected to play in gut physiology^7^ (Fig. 1i,j). Detection of ENC6 and ENC7 by their most specific markers (NMU, CCK and UCN3) was possible but suboptimal (Fig. 3), and we therefore acquired *Nmu-Cre* and *Cck-IRES-Cre* mice. Both transgenic lines were initially crossed to *R26RTomato* mice. Using RNAscope we determined that 95,7% of *Nmu*^+^ cells were labelled with TOM and that 93% of TOM^+^ cells expressed *Nmu*, confirming the efficiency and specificity of *Nmu-Cre;R26RTomato* mice (Extended Data Fig. 5a,d,e). In contrast, *Cck-IRES-Cre;R26RTomato* mice showed wide-spread TOM labelling in both enteric neurons and glia (Extended Data Fig. 5b), likely due to transient embryonic *Cck* expression^17^. To target CCK^+^ neurons, we instead injected Adeno-Associated Virus (AAV-PHP-S)^30^ carrying an inducible reporter (DIO-EYFP or DIO-Ruby3) into *Cck-IRES-Cre* mice. Reporter expression three weeks post-transduction was excluded from enteric glia and labelled *Cck*^+^ neurons (95,8% of EYFP cells; Extended Data Fig. 5c-e). To label ENC12 we chose the marker combination NTNG1/CALB identified in our immunohistochemical analysis (Fig. 3).

We first assessed the previously most used IPAN markers CGRP, NF-M and CALB protein expression as scRNA-seq indicated differential expression in the three putative IPAN classes (Fig. 1h). CGRP was observed in most ENC6 neurons, but was not readily detectable in ENC7 and 12 (Fig. 4a-d). NF-M was most prominent in ENC12 (100%), but also detected in a majority of ENC6 and ENC7 neurons. CALB (defined as marker of ENC12) was only found in a minor proportion of ENC7 and ENC6 neurons. In conclusion, previous IPAN markers formed the following combinatorial codes: ENC6 (CALB^−^ CGRP^+^ NF-M^+/−^), ENC7 (CALB^+/−^ CGRP^−^ NF-M^+/−^), and ENC12 (CALB^+^ CGRP^−^ NF-M^+^).

**Figure 4.**
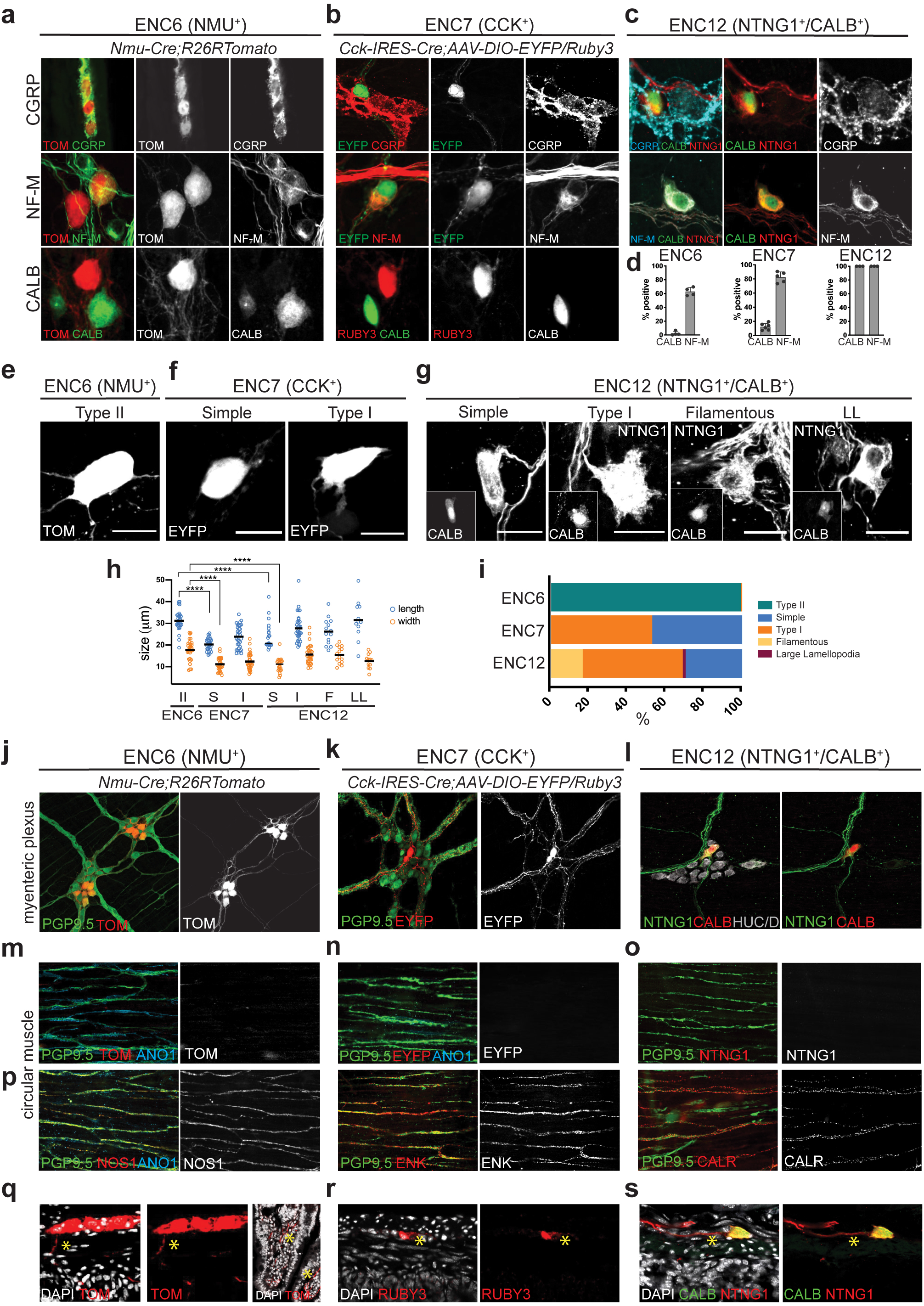
Assessment of IPAN characteristics in ENC6, 7 and 12. **a-c)** Immunohistochemical detection of IPAN markers in ENC6, 7 and 12. Pictures show either myenteric peel preparations or transverse sections. **d)** Graph showing the proportions of cells among ENC6, 7 and 12 expressing IPAN markers. Each dot indicates one animal. n= 3-6. NF-M was detected in 63,6±5,43% of ENC6, 82,5±7,42% of ENC7 and 100% of ENC12 neurons. CALB was found in 2,3±2,96% of ENC6 and 14,9±4,76% of ENC7 neurons and considered a defining marker of ENC12. We did not attempt to quanitify CGRP as faithful detection required colchicine treatment, which compromised other required marker expression. **e-g)** Representative examples of ENC6, 7 and 12 cell morphologies in *Nmu-Cre;R26RTomato, Cck-IRES-Cre;AAV-DIO-Eyfp/Ruby3* mice and wildtype mice immunohistochemically labelled with NTNG1/CALB, respectively. See Extended Data Fig. 6 for more examples. **h)** Quantification of neuron sizes in morphological groups of ENC6, 7 and 12. Each circle represents one cell. Lines indicate the mean. Student’s t-test was used for statistical analysis. ENC6: Type II ((length 31±5μm; width 18±5μm; n=30); ENC7: Type 1 (24±5μm length and 12±4μm width; n=32), Simple (20±3μm length; 11±3μm width; n=28). ENC12: Filamentous (26±6μm length; 15±4μm width; n=14), LL (32±9μm length; 13±3μm width; n=12), Type I (28±6μm length; 16±4μm width; n=33), Simple (24±6μm length; 11±3μm width; n=22) **i)** Quantification of proportions of different morphological types among ENC6, 7, and 12. Number of cells analysed: 689 ENC6 neurons from 2 animals; 440 ENC7 neurons from 2 animals; 321 ENC12 neurons from 2 animals. **j-l)** Representative pictures of cell bodies and projections originating from ENC6,7 and 12 visible in the plane of the myenteric plexus. **m-o)** Immunohistochemical analysis of ENC6, 7 and 12 projections in the circular muscle layer showing clear labeling of axons (PGP9.5^+^) and ICCs (ANO1^+^) but absence of ENC6, 7 and 12 axons in this layer **p)** Immunohistochemical detection of motor neuron projections in the circular muscle layer. Abundance of motor neuron axons (NOS1^+^, ENK^+^ or CALR^+^) contrasts with the absence of ENC6, 7 and 12 axons in this layer. **q-s)** Representative pictures of transverse sections showing the direction of projections originating from ENC6, 7 and 12 neurons. Many axons of ENC6 were found to cross the circular muscle layer and project down into submucosa and villi. Axons of ENC7 and ENC12 were only found in the plane of myenteric plexus. CALB: Calbindin; CALR: Calretinin; CCK: Cholecystokinin; CGRP: Calcitonin Gene Related Peptide; EYFP: Enhanced Yellow Fluorescent Protein; ENK: Enkephalin; HUC/D: Elav-Like Protein 3/4; NF-M: Neurofilament M; NMU: Neuromedin U; NOS1: Nitric Oxide Synthase 1; NTNG1: Netrin G1; TOM: tdTomato, DAPI: 4ʹ,6-diamidino-2-phenylindole

IPANs are thought to mainly correspond to cells with Dogiel Type II morphology, defined as large smooth and slender cell body with multiple axons.^31,32^ However, uni-axonal filamentous enteric neurons have also been attributed mechanosensory functions in the guinea pig.^33^ We labelled cellbodies/projections of ENC6 and 7 by fluorescent reporters, and ENC12 by NTNG1/CALB antibodies, to determine morphological types within these classes. ENC6 neurons were large with two or more axons and therefore classified as Type II (Fig. 4e,h,i; Extended Data Fig. 6a). ENC7 neurons were either Dogiel Type I (lamellar dendrites and one discernable axon)^31^ or what we termed “Simple”, having round or oval cell bodies and one (sometimes two) axons (Fig. 4f,h,I; Extended Data Fig. 6b). ENC12 morphologies displayed remarkable heterogeneity; we found Type I, “Simple”, Filamentous, and a population with large lamellar cytoplasmic extension, here termed “LL” (Fig. 4g-I, Extended Data Fig. 6c). We conclude that the morphology of NMU^+^ neurons (ENC6) matches previous description of Type II IPANs, while filamentous ENC12 neurons could correspond to the subset of mechanosensitive IPANs.

Given the first position of IPANs in the peristaltic circuit, they connect to interneurons and motor neurons within the myenteric plexus but do not innervate the external muscle layers, where motor neuron projections predominate. While axons from the three ENCs were readily observable in the plane of the myenteric plexus (Fig. 4j-l), none were identified within the circular muscle (Fig. 4m-o). In contrast, extensive staining for inhibitory (NOS1^+^) and excitatory (ENK^+^, CALR^+^) motor neurons were found in the axonal tracts (PGP9.5^+^), that together with Interstitial Cells of Cajal (ANO1^+^) parallel the axis of the circular muscle (Fig. 4p). Thus, we concluded that ENC 6, 7, and 12 are not motor neurons.

Another distinguishing feature of IPANs is their projection across the muscle layer to the submucosa and mucosa. On transverse tissue sections, we found ENC6 axons intersecting the circular muscle and in the villi (Fig. 4q). In contrast, axons from ENC7 and ENC12 were only observed in the plane of the myenteric plexus and not in the muscle or villi (Fig. 4r).

In conclusion, our study identified ENC6 as Type II IPANs and with that the non-ambiguous marker NMU and the *Nmu-Cre* mouse strain as a useful tool to further study and modulate this important enteric neuron type. Expression of serotonin receptors (e.g. *Htr3a,b*, Extended Data Fig. 3a) make ENC6 equipped to respond to mucosal serotonin-producing enterochromaffin cells.^3^ The particular morphology displayed by a subset of ENC12 (NTNG1/CALB) neurons and the expression of PIEZO2 (Figs. 2b, 3) connects these cells to previously reported filamentous IPANs. Our analysis indicate that ENC7 (CCK neurons) do not correspond to *bona fide* IPANs, despite their expression of NF-M and CALB. Thus, common IPAN markers cannot faithfully discriminate IPANs from other neuron types.

### Single Cell RNA-sequencing reveals transcriptional programs of generic cell states of the developing ENS

To understand the generic and specific differentiation processes underlying the formation of ENC1-12 we performed scRNA-seq of the whole ENS isolated from *Wnt1Cre-R26RTomato* small intestine at E15.5 and E18.5, when most myenteric neuron types have been/are generated.^15,16^

A total of ∼3000-3500 transcriptomes were captured per stage. After removal of unhealthy, ambiguous or non-ENS cells, the remaining cells (3260 for E15.5 and 2733 for E18.5) were subjected to downstream analysis. An overall similar manifold was observed at both stages (Fig. 5a,b). The enteric progenitor/glial marker *Sox10* (Fig. 5c) was used to assign clusters corresponding to progenitor cells within different parts of the cell cycle (Fig. 5d), while *Elavl4^HIGH^* expression (Fig. 5c) demarcated differentiating neurons. A clear *Ascl1*^HIGH^ cluster at the intersection of the two was considered to correspond to cells undergoing neurogenesis (here termed neuroblasts). While glial differentiation occurs earlier^34^, we only observed clear clusters corresponding to glia (*Plp1*^high^; S100b^high^) at E18.5 (Fig. 5b). The two populations appeared to be the previously reported Schwann Cell Precursors (SCP) that and contribute to the ENS at perinatal stages^13^ (SCP markers: *Dhh*, *Mal* and *Mpz;* mousebrain.org), and vagal neural crest-derived enteric glia (enteric glia markers: *Apoe, Nkain4;* mousebrain.org).

**Figure 5.**
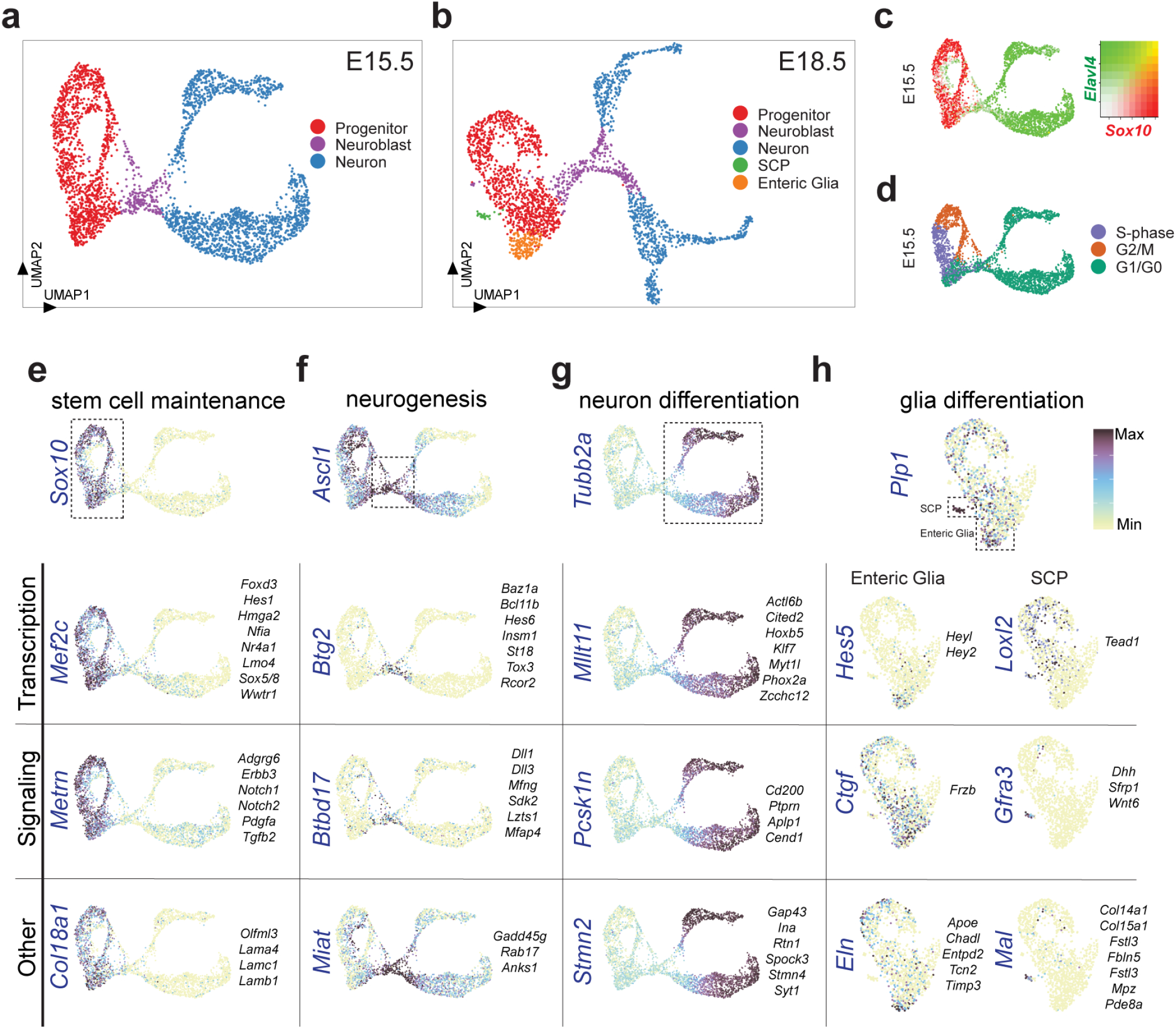
Single cell transcriptome analysis of the developing ENS reveals transcriptional programs of generic cell states. UMAP representation of E15.5 **(a)** and E18.5 **(b)** ENS scRNA-seq datasets displaying generic states of the developing ENS. **c)** Complementary expression of *Sox10* and *Elavl4* reveals progenitor/glia cells versus neuronal populations in UMAP. Color map represents scaled expression of *Sox10* (red), *Elavl4* (green), co-expressed (yellow) and non-expressing (grey). **d)** Cell cycle assignment mapping onto UMAP at E15.5. Genes organized into categories, with likely functions in stem cell maintenance (E15.5; **e**), neurogenesis (E15.5; **f**), neuron differentiation (E15.5; **g**) and two glial populations (E18.5; **h**) plotted on UMAPs. Color bars indicate expression level with maximum cut off at the 90^th^ percentile. UMAP: Uniform Manifold Approximation and Projection; ENC: Enteric Neuron Class

Differential expression analysis (Extended Data Fig. 7) of these broad cell states discovered transcription and signaling genes with likely functions in stem cell maintenance (*Mef2c*, *Sox5/8, Notch1/2*; Fig. 5e), neurogenesis (*Dll1*, *Dll3*, *Hes6*; Fig. 5f), neuron differentiation (*Myt1l*, *Ptprn*; Fig. 5g), SCP (*Gfra3*, *Wnt6;* Fig. 5h) and enteric glia (*Hes5*, *Frzb;* Fig. 5h). The Notch signaling pathway appeared strongly implicated in regulating basic ENS development given the differential expression of its molecular components across the cell states (Extended Data Fig. 8). Neuroblasts displayed enriched expression of various chromatin modifiers (*Baz1a*, *Tox3*, *Bcl11b*) likely to execute the vast transcriptional change associated with the progenitor-to-neuron differentiation.

### Enteric Neuron Classes are formed through a binary fate split at neurogenesis followed by successive diversification at the post-mitotic stage

UMAP (Uniform Manifold Approximation and Projection) representations of ENS at E15.5 and E18.5 show that enteric neurons are formed through two trajectories, which we term Branch1 and Branch2. On these trajectories, we sought to locate cells that have acquired traits of ENC1-12. We first projected principal component analysis (PCA) structure of the juvenile scRNA-seq data onto the developmental data and, within this space, searched for anchors based on shared overlap of mutual neighbors. The transfer of class identities was achieved by a weigh vote classifier, giving a prediction score for each ENC on the developmental data (cells with high score received consistent votes across anchors and were mapped with high confidence). Mapping of transferred identities with predicted score >0.5 onto the trajectories indicated that ENC1-7 are generated through Branch1, while ENC8-12 differentiate in Branch2 (Fig. 6a,b). Selective marker expression further confirmed traits of ENC1/2 (*Ndufa4l2*) and ENC8/9 (*Nos1, Gal, Vip*) within the early branches (Fig. 6c). Branch1 diverged further into Branch1a, corresponding to emergence of ENC3/4 (*Gda, Penk, Fut9*) or Branch1b, corresponding to ENC6 (*Ano2*) and ENC7 (*Mgat4c, Sema5a, Ucn3*). Within Branch2, traits corresponding to ENC10 (*Neurod6*) and ENC12 (*Ntng1, Calb1, Nxph2*) gradually appeared (Fig. 6a-c). ENC5 and ENC11 neurons must emerge after E18.5 as few or no cells were mapped to these identities. Notably, genes conferring class-specific functional features were already apparent including *Oprk1* (ENC1-3), *Gucy2g* (ENC4), *Grin3a* (ENC6) and *Piezo2* (ENC12) (Fig. 6d; compare with Fig. 2a-c). ENC10-12 identities coincided with a nitrergic-to-cholinergic switch (*Nos1* down- and *Slc18a2* up-regulated; Fig. 6c). The mixed nitrergic/cholinergic identity of the mature ENC10 (Fig. 1h) could therefore be explained by their generation at the intersection of this process.

**Figure 6.**
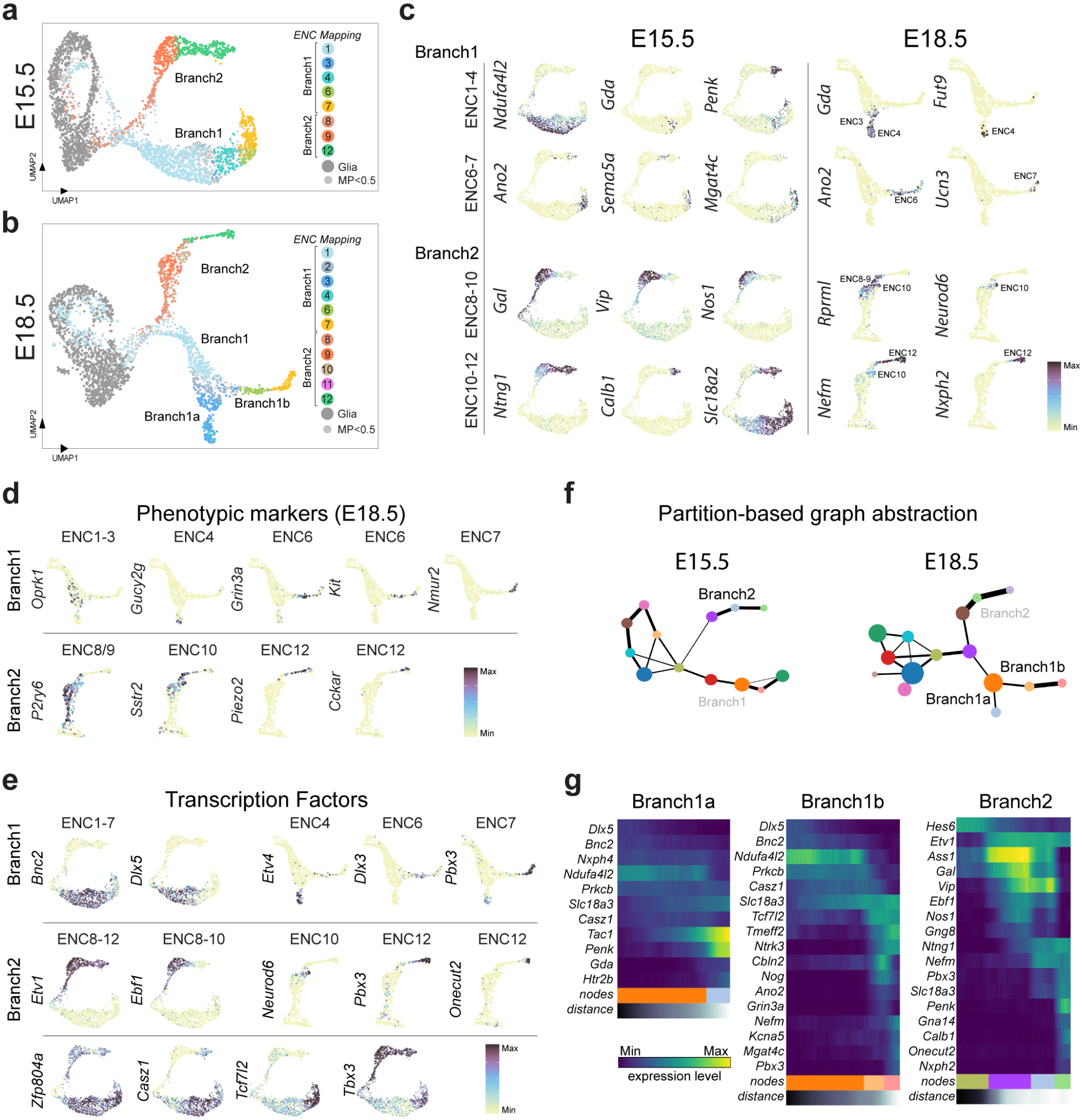
Enteric Neuron Classes emerge through an initial binary split followed by post-mitotic neuron diversification. **a,b)** UMAP representation of E15.5 and 18.5 data sets with identity transfer of ENC1-12 and ‘enteric glia‘. Most progenitor cells mapped to enteric glia as due to their overall similar gene expression. Threshold for maximum prediction score was set to > 0.5;cells. Small grey dots indicate cells with no identity transfer in this range. **c)** ENC1-12 juvenile marker genes displayed on E15.5 and E18.5 feature plots validates ENC assignments. **d)** Phenotypic markers displayed on UMAP graphs at E18.5. **e)** Transcription factors displayed on feature plots including those limited to each Branch, selective ENCs or groups of ENCs. Most expression patterns appear to remain in the juvenile ENCs (compare with Fig 2d). Color bars indicate expression level with maximum cut off at the 90^th^ percentile. **f)** PAGA graphs on UMAPs of E15.5 and E18.5 ENS corresponding to the differentiation of ENC1-4 (Branch1a), ENC6-7 (Branch1b) and ENC8-12 (Branch2). Weighted edges represent degree of significant connectivity between partitions. Node coloring is arbitrary. **g)** Heatmaps organized by diffusion pseudotime indicating gradual gene expression changes in the three PAGA paths. UMAP: Uniform Manifold Approximation and Projection; ENC: Enteric Neuron Class; PAGA: Partition-based graph abstraction

We screened for transcription factors that define branches and their differentiation into ENCs. The early segregation between branches at E15.5 was associated with *Bnc2/Dlx5* (Branch1), or *Etv1/Ebf1* (Branch2) (Fig. 6e). Thus, the division between ENC1-7 and ENC8-12 by the complementary expression of *Etv1* and *Bnc2* (see Fig. 2e) indeed reflects an early binary branching process. *Etv4* demarcated Branch 1a (ENC4), while *Dlx3* was found within Branch1b (ENC6). *Neurod6* (ENC10) appeared further down Branch2 and was followed by *Pbx3* and *Onecut2* (ENC12) (Fig. 6e). The onset of most transcription factors appeared to reflect emergence of specific ENCs and remained until juvenile stages (see Fig. 2d), while others displayed a broad developmental expression (*Zfp804a*, *Ebf1*). *Etv4* and *Dlx5* were not detected in mature ENCs, indicating functions limited to early diversification and not cell identity maintenance.

To investigate gene expression along developmental trajectories, we defined partition-based graph abstraction (PAGA) representation of E15.5 and E18.5 data. PAGA coupled with diffusion pseudotime analysis confirmed the gradual differentiation of ENCs within Branch1a, Branc1b and Branch2 (Fig. 6e). Thus, the 12 ENCs appear not to be generated from 12 different stem cell pools. Instead, proliferating progenitors undertake only two prototypic traits associated with ENC1 or ENC8 (excitatory versus inhibitory motor neurons) upon completing neurogenesis. Subsequently, many neurons mature within their first identity, while others downregulate their specific markers and convert into new classes (ENC2-7; ENC9-12).

### The ENC8/9 to ENC12 phenotypic switch depends on PBX3 expression

As NOS1^+^ neurons are of particular clinical importance^8,9^, we decided to focus on the gradual diversification within Branch2. We noted a border between ENC8/9 and ENC12 (Fig. 6a,g), coinciding with opposing expression of *Nos1/Gal/Vip* relative to *Ntng1/Calb1/Nfem* (Fig. 6c). This indicates that neurons obtaining early traits of ENC8/9 convert into ENC12 neurons. Notably, *Pbx3* expression correlated with the transition point (Figs. 6g, 7a). We have previously shown that PBX3 is expressed in enteric neurons of both mouse and human embryonic gut and is associated with CALB^+^, but not NOS1^+^ expression.^17^ Quantification at E18.5 revealed that only 1.5±0.4% (n=3) of the NOS1^+^ neurons expressed PBX3, and those neurons displayed low levels of NOS1., The expression dynamics suggests that the ENC8/9 identity may be negatively regulated by PBX3.

**Figure 7.**
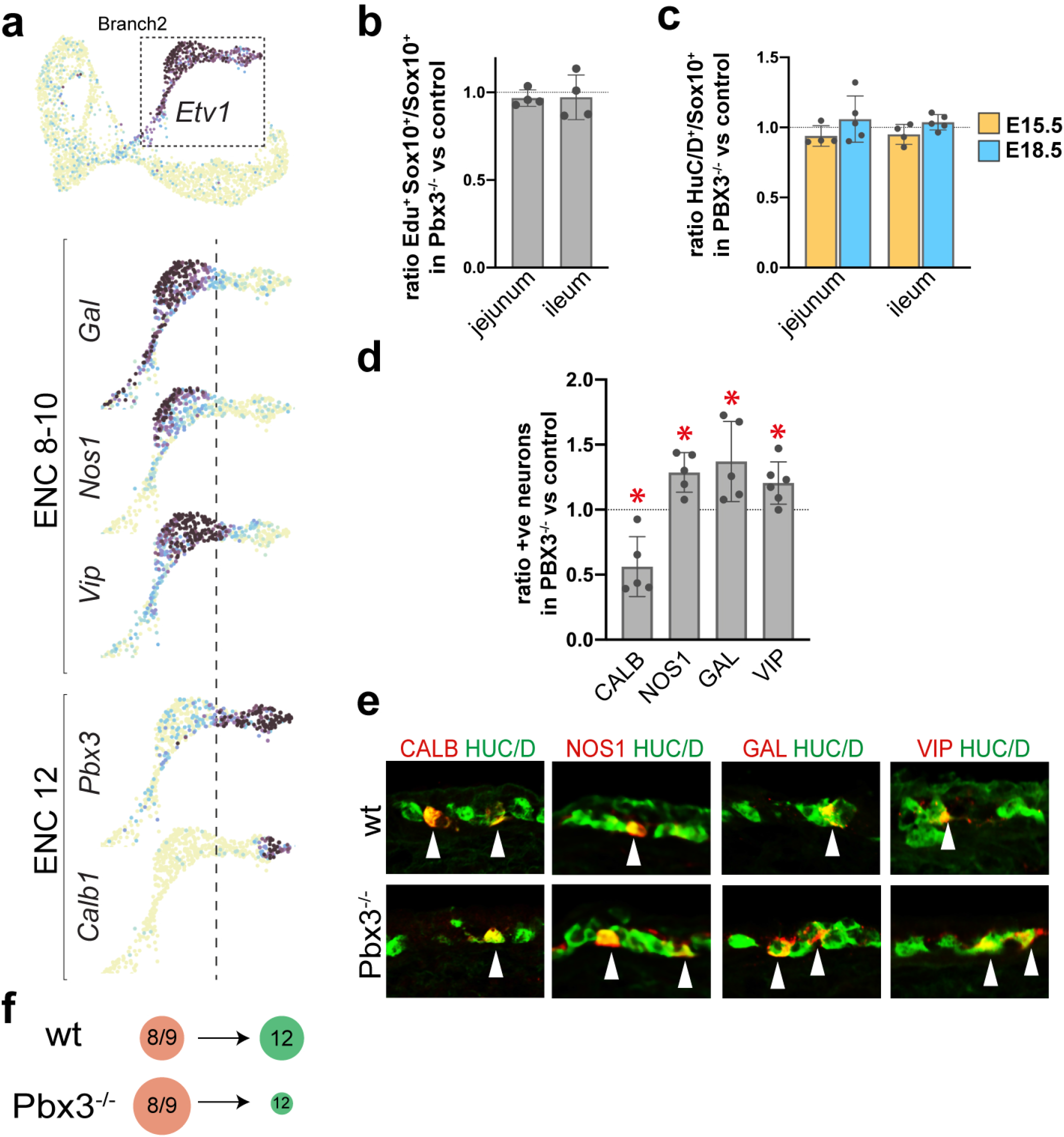
Loss of PBX3 expression impairs the conversion of ENC8/9 to ENC12 traits. **a)** Feature plots showing the expression profile change in Branch2 (Etv1^+^) at E15.5. Dotted line demarcates the approximate border between ENC8/9 and ENC12 and coincide with onset of *Pbx3* expression. **b)** Graph showing that the ratio of SOX10**^+^** cells that incorporated EdU after a 90 min pulse injection is similar in *Pbx3^−/-^* mutants and *wt* controls (set to 1) at E15.5. n = 4. **c)** Graph showing an unchanged ratio of HUC/D^+^ over SOX10^+^ cells in *Pbx3^−/-^* mutants compared with *wt* controls (set to 1) at E15.5 and E18.5. n = 4-5. **d)** Graph showing the ratio between the percentages of neurons expressing different neurotransmitters/peptides in *Pbx3^−/-^* mutant and control (set to 1). n = 5-6. *p < 0.05. **e)** Representative pictures of the small intestine at E18.5 in *wt* and *Pbx3^−/-^* embryos showing increased ratios of neurons expressing NOS1, GAL and VIP and decreased ratio of neurons expressing CALB**. f)** Schematic drawing indicating the increased number of presumed ENC8/9 at the expense of ENC12 neurons in the ENS of *Pbx3^−/-^* mutant mice compared to control. Student’s Paired t-test used in (d). wt: wildtype; EdU: 5-ethynyl-2 deoxyuridine; HUC/D: Elav-Like Protein 3/4; GAL: Galanin: NOS1: Nitric Oxide Synthetase 1; CALB: Calbindin; VIP: Vasoactive Intestinal Peptide.

To evaluate the role of Pbx3 in ENS development, we analyzed the guts of *Pbx3^−/-^* mutant mice. As *Pbx3^−/-^* mutant mice die within hours after birth due to central respiratory failure^35^ we focused our analysis at E15.5-E18.5. We first investigated if the proliferative capacity of ENS cells was affected in *Pbx3^−/-^* mutant mice. The ratio of SOX10^+^ cells in the S-phase of the cell cycle, incorporating the nucleoside analog EdU after a 90 min pulse, was the same in mutant and control guts at E15.5 (Fig. 7b). Moreover, the ratio of HUC/D^+^ cells to SOX10^+^ cells was unchanged in the mutant compared to control guts (Fig. 7c). Thus, loss of *Pbx3* does not impact generic programs of enteric proliferation or neurogenesis.

The correlation between *Pbx3* expression in emerging ENC12 neurons bordering ENC8/9 neurons prompted us to determine the percentage of HUC/D^+^ neurons expressing the ENC8/9 markers GAL, NOS1 and VIP and the ENC12 marker CALB at E18.5. We found a significant increase in the proportion of neurons expressing VIP (20.5%), NOS1 (28.7%) and GAL (37.1%) (Fig. 7d,e), while that of CALB1 neurons was decreased by 43.8% (Fig. 7d,e) in *Pbx3^−/-^* mutants compared to control guts. Thus, our data suggest that PBX3 suppresses the ENC8/9 phenotype and allows or induces differentiation of the ENC12 identity. In the absence of PBX3, NOS1^+^/GAL^+^/VIP^+^ neurons fail to differentiate efficiently into CALB^+^ ENC12 neurons (Fig. 7f).

## DISCUSSION

The neuronal composition of murine ENS and how it emerges during development have not been determined in detail. This study provides a new taxonomy of small intestine myenteric neurons and a novel principle of their embryonic diversification.

### A novel definition of myenteric neuron classes in the mouse small intestine

We present a pan-neuronal mouse line, Baf53b-Cre, that allow specific and efficient targeting of ENS neurons, a tool which has been lacking in the ENS field. We define 12 subclasses of myenteric ENS neurons, (ENCs) and suggest their basic functional assignments (IPAN, motor neuron and interneuron). Although previously used IPAN markers are found within three ENCs, our characterization of morphology and projections suggest that IPANs may only correspond to ENC6 and subsets of ENC12 neurons. Thus, our validated genetic mouse tool (*Nmu-Cre*) and stringent markers (NTNG1/CALB) together with their communication profiles (Extended Data Fig. 3) should help future interrogation of sensory modalities within the ENS. We do not rule out that other ENCs (or subsets within) form additional IPAN types, including ENC7. A plausible alternative role of ENC7 are viscerofugal as CCK expression has been associated with this neuron type in the guinea-pig.^3^ However, the relative abundance of ENC7 cells along the small intestine suggest they also sub-serve other functions, which should be possible to delineate by selective manipulation of ENC7 using *Cck-IRES-Cre* mice. Our molecular classification of motor neurons are in agreement with the previous major division into inhibitory and excitatory types^3^, and we also identified plausible subtypes (ENC1-3 and ENC8-9, respectively). ENC4 and 11 represented rare neuron types with highly unique expression patterns, including a plausible novel neurotransmitter (noradrenaline). ENC5 and ENC10 were assigned as interneurons, but knowledge of these neuron types is limited. Thus, further functional dissections of these four classes are warranted, which could be aided by already commercially available transgenic mice targeting for instance Nfatc1, Dlk1, Sst and Neurod6.

While the majority of differentially expressed genes was associated with classical neuron functions (neurotransmitters, ion channels, signaling factors), we uncovered a set of unexpected marker genes. NDUFA4L2, a mitochondrial reprogramming factor under hypoxic conditions ^36^, was exclusively expressed in developing and juvenile ENC1-3 and has not been detected in neurons of other nervous systems (mousebrain.org; data not shown). RPRML, a protein of unknown function is unique to NOS1^+^ neurons (ENC8-10) in the peripheral nervous system (mousebrain.org; data not shown). Thus, the unprecedented level of detail within our novel ENS classification may open up new research questions, offering a rich resource for basic and translational research.

### A novel model for cell diversification in the developing ENS

Although cell proliferation, migration and survival are well understood processes within the developing ENS ^12^, neuronal diversification has received little attention.^17,37^ In contrast, gene regulatory programs driving neuronal subtype differentiation in the developing CNS have been comprehensively resolved.^38^ This lag of knowledge in ENS development may stem from the fundamentally different ways in which these nervous systems are formed. In the CNS, the process from committed progenitor to specified neuron subtype occurs at stereotypical positions within the neural tube, making studies of gene regulatory networks attainable.^14^ In the developing gut, progenitor cells and differentiating neurons are intermingled in seemingly stochastic manners, and the motile behavior of progenitors^39^ is likely to perturb morphogens to set up stable differentiation programs based on their position. Instead, a temporal mechanism has been the suggested dominant mode of specification within the developing ENS. In support, birth-dating studies signify that the neurogenic peak of phenotypically different neurons occurs at different developmental timepoints.^15,16^ The present study brings forward a new model of enteric neuron specification. We show that primitive features of only two subsets of enteric neuron subclasses (ENC1 and ENC8) are generated as dividing progenitors undergo neurogenesis. Subsequent identity conversions appear to underlie formation of ENC2-7 and ENC9-12. Thus, the 12 ENCs are generated from an initial binary difference, which become further diversified at the post-mitotic state. In support, the expression of CGRP and 5-HT, which we here map to ENC6 and ENC12 (Figs. 4a, 3) is turned on 4-6 days (E17-18)^37^ after the birth dates of these neurons (neurogenic peaks at E11.5 and E13, respectively)^16^. Furthermore, retrograde labelling at E11.5 to E16.5 showed that all neurons had single long processes, indicating that the Dogiel Type II neurons (which we identify as ENC6) develop at later stages.^40^ Thus, the developing ENS could rely on flexible postnatal neurons instead of niches for heterogeneous stem cell populations. This principle might explain how the ENS can be formed without spatially defined stem cell populations and during a protracted time (until P30).^41^ The two major branch identities must however already be segregated in progenitors undergoing their last cell-cycle, as a previous Confetti-based lineage-tracing study showed that clones consisting of 2-4 cells, traced at E12.5 and analyzed in adult mice, never contained both NOS1^+^ (ENC8-10) and CALR^+^ (ENC1-2) neurons.^34^ Our data indicates that the subsequent conversions of ENC1/2 or ENC8/9 traits are associated with switching of a number of critical phenotypic genes. As part of the ENC8/9 to ENC12 conversion, *Nos1/Gal/Vip* is downregulated and *Calb1* is upregulated, a process critically dependent on the activity of the transcription factor PBX3. Transient neuronal expression of NOS1 is supported by an earlier study which determined the birth-wave of NOS1^+^ neurons to commence at E12.5 although NOS1^+^ neurons were readily observed already at E11.5.^16^ Thus, although ENC8/9 and ENC12 are generated from the same birth branch, ENC12 could represent neurons that are born at early developmental time points and switch identity, while later born neurons retain their ENC8/9 identity. Such model would make our data consistent with studies showing a correlation between birth date and cell identity.^15,16^ We have previously assembled a gene expression resource of the developing ENS.^17^ Surprising to us, the greatest diversity of plausible cell identity regulatory genes (transcription and signaling factors) was found amongst neurons and not within the dividing progenitor population. Thus, also this study supports a sequential differentiation process within already post-mitotic neurons as major mode of diversification within the ENS.

### Generic differentiation of enteric neurons and glia

We identified transcriptional profiles representing each of the generic states of dividing progenitors, neurogenic cells (neuroblasts), differentiating neurons and two glial types. Many progenitor state transcription factors (including *Mef2c* and *Sox5*) were also enriched in a microarray comparing SOX10^+^ cells to the whole developing ENS^17^ and scRNA-seq of ENS at E13^34^ further supporting their role in stem cell regulation. Neurogenic cells assembled into a single tight cluster, indicating a strong generic neurogenesis program. The transcriptional code resembled to a large extent that of differentiating adrenergic chromaffin cells, also expressing for instance *Dll3, Hes6, Gadd45g*.^42^ Further cross-comparisons to other neural-crest related lineages could determine the generic neural crest transcriptional core versus ENS-specific transcriptional networks.

We present the transcriptional profile of enteric SCP.^13^ While our analysis is unable to specifically reveal SCP-derived cell differentiation trajectories we identified genes (*Mpz* or *Mal*) that may be used as basis to design transgenic lineage tracing. Our data also include generic traits of basic ENS cellular components (progenitor/glia/neuron) that may regulate the organization of the ENS. For instance, phenotypic genes in glial/progenitor populations were predominated by extracellular matrix molecules (e.g. collagens, laminins, *Eln*, *Mal;* Fig 5e-h), which possibly could support accurate neuron differentiation, survival and network formation.

### Impact on future regenerative medicine for enteric neuropathies

Several primary enteric neuropathies have been shown to mainly affect enteric NOS1^+^ neurons including esophageal achalasia and hypertrophic pyloric stenosis. Enteric NOS1^+^ neurons are also damaged in Chaga’s disease and diabetic gastroparesis.^8,9^ NOS1 neurons may be especially susceptible to damage due to their production of free radical nitric oxide (NO), which could react with reactive oxygen species created in stressed tissue.^9^ Esophageal achalasia has been pointed out as plausible first disease for novel cell-based therapies as its neural deficit is well defined and localized within a small tissue area.^11^ Engineering of NOS1 neurons, whether *in vivo* or *in vitro* requires detailed knowledge of NOS1 neuron differentiation. Our study describes the generation of NOS1^+^ motor neurons (ENC8/9), provides early and specific marker genes and find a negative transcriptional regulator (*Pbx3*). We also present the generic neurogenic transcriptional core, knowledge needed in efforts to convert enteric glia to neurons. With further functional interrogation of the herein identified transcription factors, selective production of NOS1^+^ neurons is likely conceivable following the same principle used to make various types of brain neurons^43,44^.

## Acknowledgements

Cell sampling was performed at the Eukaryotic Single-cell Genomics core facility at Science for Life Laboratory, Sweden. Sequencing services were provided by the National Genomics Infrastructure at Science for Life Laboratory, Sweden. U.M was supported by The Swedish Research Council (Vetenskapsrådet; 2016-03130), Swedish Medical Society (SLS), Ruth and Richard Julin Foundation, Ollie and Elof Ericssons Foundation, Magnus Bergvall Foundation and Åke Wiberg Foundation. A.M was supported by Wenner-Gren Foundations. F.M was supported by the Brain Foundation (Hjärnfonden). P.E acknowledges ERC (PainCells 740491), The Swedish Research Council, KAW Scholar and project grant, and Welcome Trust (200183).

## Author Contributions

Study concept and design: A.M, K.M, V.K and U.M

Acquisition of data: A.M, F.M, K.M, V.K, R.K, W.L and U.M

Analysis and interpretation of data: A.M, F.M, K.M, V.K, R.K, W.L, and U.M

Drafting of Manuscript: A.M, K.M, V.K, P.E and U.M

Obtained Funding: F.M, P.E and U.M

## MATERIAL & METHODS

### Mice

The generation of the *Baf53b-Cre*^1^ (JAX, #027826), *Cck-Ires-Cre*^2^ (JAX, #12706), *Nmu-Cre* (MMMRC, 036643-UCD), and *Wnt1-Cre*^3^ mouse strains have been previously described. Strains were crossed with *Ai14*^4^ *(R26RTomato;* JAX #007908). *Pbx3^+/−^* mice were kindly provided by C. Villaescusa (Karolinska Institutet) and have been described^5^. *Pbx3^+/−^* and *Baf53b-Cre* mice were kept on a C57BL/6 background. Animals were group-housed, with food and water *ad libitum,* under 12-h light-dark cycle conditions. Animal experiments were approved by the local ethics committee in northern Stockholm (N87/13 and 5264/18).

### Tissue preparation for single cell RNA sequencing

#### Juvenile tissue preparation

A total of four *Baf53b-Cre*;*R26RTomato* mice, postnatal day (P)21, were used for the single cell experiment. During all stages of the dissociation protocol, the tissue was kept in dissection solution: TRIS-HEPES recovery solution containing 76 nM Tris HCl, 19.5 mM Tris Base, 2.5 mM KCl, 1.2 mM NaH_2_PO_4_, 30 mM NaHCO_3_, 20 mM glucose, 5 mM sodium ascorbate, 3 mM sodium pyruvate, 0.5 mM CaCl_2_ and 10 MgSO_4_ (pH 7.3-7.4). The dissection solution was equilibrated in 95% O_2_, 5% CO_2_ for 30 min before the start of the experiment and held on ice at all steps. The dissected small intestines were cut in 5 cm pieces and each segment flushed inside with ice-cold dissection solution. The mesentery was removed and intestine pieces opened lengthwise along the mesenteric border. The tissue pieces were pinned with the mucosa side down on a Sylgaard (Dow Corning) covered dissection dish on top of an ice-block. The smooth muscle layers, containing the myenteric plexus were peeled off from the submucosa using watchmaker forceps and stored in dissection solution on ice until all pieces were peeled. Peels were cut in 2-5 mm^2^ pieces and placed in 5 ml digestion solution (0.75 mg/ml Liberase TH Research grade (Roche), 0.1 mg/ml DNAseI (Sigma-Aldrich) and 25U/ml Dispase (Corning)) in DMEM/F12 at 37°C for 30 minutes, with gentle shaking of the tube every 10 min. After completed digestion, enzyme mixture was replaced with 2.5% bovine serum albumin (BSA; Sigma-Aldrich), 5mM EDTA in ice-cold DMEM/F12. Tissue pieces were triturated using three fire-polished Pasteur pipettes with decreasing opening size (from ∼70% to 10% of the original opening size) that had been coated with 1% BSA solution, and then filtered through a DMEM/F12-equilibrated 30 μm cell strainer (Miltenyi Biotec). Single cells were collected by centrifugation at 120g for 5 min at 4°C. The pellet was resuspended in 2 ml DPBS solution, 1% BSA and 1% DRAQ7 (Biostatus). Tomato^+^ cells were sorted by flow cytometry on a BD Influx™ cell sorter and collected in ice-old PBS containing 0.04% BSA.

#### Embryonic tissue preparation

Embryonic day (E) 0,5 was considered when vaginal plug was seen in female mice. *Wnt1-Cre*;*R26Tomato* were used for single cell RNA sequencing experiments. In total 5 and 3 embryos at E15.5 and E18.5 respectively, were used and for each developmental stage the small intestines were pooled. During all stages of the dissociation protocol the tissue was kept in DMEM/F12 medium (Invitrogen) on ice, except where mentioned otherwise. Dissection steps were carried out under a stereo microscope. The mesentery was removed, the small intestines were cut into 2-5mm^2^ pieces and put into digestion solution (0.75mg/ml Liberase TH Research Grade (Roche), 12 U/ml Dispase, 0.1mg/ml DNAseI (Sigma) in DMEM/F12) at 37°C for 10 min (E15.5) or 20 min (E18.5) with shaking every 5 min. The enzyme mixture was replaced with DMEM/F12 medium containing 2% BSA and 5mM EDTA. The cells were then manually triturated using 3 fire-polished Pasteur pipettes with decreasing opening size (from ∼70% to 10% of the original opening size) that were previously coated with 1% BSA solution for at least 1h at RT. The single cell suspension was filtered through a DMEM/F12-equilibrated 30μm cell strainer (Miltenyi Biotec) and centrifuged at 120g for 5 min at 4°C. Cells were resuspended in DPBS containing 1% BSA and 1% DRAQ7 (Biostatus). Tomato+ cells were sorted on a BD Influx™ cell sorter, collected in ice cold PBS containing 0.04% BSA and passed on for single cell RNA sequencing analysis.

### Single cell RNA sequencing

The sampling was carried out with 10x Genomics Chromiusm Single Cell Kit Version 2 (10x Genomics, CA, USA) at Eukaryotic Single Cell Genomics (ESCG), Stockholm, Sweden. Cell suspensions were adjusted to 500-1000 cells/μl, and added to 10x Chromium RT mix to achieve loading target numbers between 2500-8000 cells. The manufacturer’s instructions were followed for downstream cDNA synthesis, library preparation, and sequencing using Novaseq-S1 (Illumina).

### Virally-mediated neuron labeling

ssAAV-PHP.S/2-shortCAG-dlox-EYFP(rev)-dlox-WPRE-hGHp(A) (abbreviated as PHP.S-DIO-EYFP) and ssAAV-PHP.S/2-shortCAG-dlox-mRuby3(rev)-dlox-WPRE-hGHp(A) (abbreviated as PHP.S-DIO-mRuby3) were constructed at the Viral Vector Facility (VVF) at the University of Zurich (# v309-PHP.S and v378-PHP.S respectively) using Addgene plasmids pUCmini-iCAP-PHP.S (#103006) and pAAV-CAG-DIO-EYFP (#104052). Procedure was performed as described.^6^ *Cck-Ires-Cre* and *Nmu-Cre* mice were injected intravenously into the tail vein with 5.7*10^11^ viral dose. Animals were sacrificed 2-3 weeks after injection and tissue analyzed.

### Tissue preparation for immunofluorescence (IF) analysis

#### Juvenile and adult small intestine sections

P21-90 small intestine was dissected, mesentery removed, and duodenum, jejunum and ileum separated. Intestinal pieces were flushed clean with ice-cold PBS and fixed overnight in 4% paraformaldehyde (PFA) in PBS at 4°C. Some tissue was prepared as swiss-rolls.^7^ Briefly, the opened tissue pieces were placed flat, mucosa side down on the bottom of an empty petri dish. A toothpick was placed at one end and the tissue was rolled up to form a roll. The rolls were pinned to a Sylgaard-covered dissection dish and fixed in 4% PFA at 4°C over-night. Tissues were then washed three times with PBS and incubated in 30% sucrose in PBS overnight at 4 °C and embedded in O.C.T. (Histolab, Leiden, The Netherlands), snap frozen and stored at −80 °C. The tissue was cut at 14 μm (20 μm for RNAscope^®^) and kept at −20 °C until use.

#### Adult myenteric plexus peels

Duodenum, jejunum and ileum intestinal segments from P21-90 mice were cleaned of mesentery and opened lengthwise along the mesenteric border. The intestinal contents were rinsed out with ice-cold PBS. The tissue was stretched and pinned with the mucosa side down on a Sylgaard-covered dissection dish and fixed in 4% PFA at 4°C overnight. Tissue was then washed three times with PBS, longitudinal muscle-myenteric plexus-circular muscle layer peeled off (myenteric peels), cut into 1 cm^2^ segments and stained directly or stored in 100% methanol at −20°C. In some cases, stretched fresh tissue was cultured overnight in DMEM (ThermoFisher) containing 10% heat-inactivated fetal bovine serum (ThermoFisher), 100 U/ml penicillin (ThermoFisher), 100 ug/ml streptomycin (ThermoFisher) and 0.1 mg/ml colchicine (Sigma). Thereafter, myenteric peels were prepared as above.

#### Embryonic tissue

Guts from E15.5 and E18.5 embryos were dissected out and fixed in 4% PFA in PBS at 4°C for 2h. Samples were subsequently washed in PBS and cryoprotected in 30% sucrose in PBS at 4°C overnight. The tissue was then embedded in O.C.T. and frozen at −80°C. Samples were sectioned at 12-14 μm, dried at RT (room temperature) for 1h and further processed for IF or frozen at −20°C.

### Immunofluorescence analysis

Frozen tissue sections (juvenile and embryonic) were air dried at RT for 1 hour, rinsed with PBS and incubated in blocking solution containing purified donkey anti-mouse Fab (Jackson, #715-007-003) diluted 1:50 in PBS for 1 hour at RT. Sections were then blocked with 2% normal donkey serum (NDS, Jackson) and 0,1% Triton X-100 (Sigma) in PBS for 1 hour and incubated with primary antibodies diluted in the same solution overnight at 4°C. They were then washed 3 times with PBS (10 minutes each) and incubated with secondary antibodies at RT for 1 hour. Tissue was washed 3 times with PBS (10 minutes each) and mounted in DAKO mounting medium (Agilent) containing DAPI.

Myenteric peels were incubated in blocking solution containing purified donkey anti-mouse Fab diluted 1:50 in PBS and 0,3% Triton X-100 overnight at 4°C. The blocking solution was replaced by the incubation buffer containing 2% NDS, 1% BSA and 0,3% Triton X-100 in PBS for 8 hours at 4°C. The tissue was incubated in primary antibodies diluted in the incubation buffer for 48 hours 4°C, washed 3 times with PBS (30 minutes each) and then placed in the secondary antibodies diluted in the incubation buffer for 2 hours at RT. The tissue was washed 3 times with PBS (15 minutes each) and mounted on glass slides using DAKO mounting medium containing DAPI.

For a complete list of primary and secondary antibodies, see Supplementary Table 2.

### EdU labeling

Time plug-mated *Pbx3^−/-^* mice received a single intraperitoneal injection of 5-ethynyl-2’-deoxyuridine (EdU; 0,1mg/g animal; Invitrogen) at E15.5. Mice were sacrificed 90 min after injection. Embryos were prepared for immunohistochemistry as described above. Tissue sections were first immuno-stained for SOX10 and then reacted for EdU using the Click iT^®^ EdU Alexa Fluor ® 488 Imaging kit (Invitrogen). After 20 minutes slides were washed in PBS and mounted.

### RNAscope^®^

RNAscope^®^ was performed according to manufacturer’s instructions (RNAscope^®^ Multiplex Fluorescent Reagent Kit v2 Assay). Protease III was used for tissue digestion. The following probes were used for the analysis: Mm-Nmu (Ref 446831) and Mm-Cck-C2 (Ref 402271-C2). Expression of fluorescent reporter proteins was evaluated by IF after the RNAscope^®^ procedure as described below.

### Imaging

Imaging of single plane or z-stacks was performed using Zeiss LSM700 or LSM800 confocal microscopes (Oberkochen, Germany), and processed in Fiji image analysis software (National Institutes of Health, Bethesda, MD).

### Cell counting and statistical analysis

Number of cells expressing various markers in the myenteric peel preparations were counted from 1 cm^2^ pieces of both jejunum and ileum in at least 3 animals under a Zeiss fluorescent microscope. Percent expression was calculated on the combined jejunum and ileum counts. Student’s *t* test was used for statistical analysis. For RNA Scope, number of Reporter^+^ / RNA^+^ cells was evaluated in 20μm sections from jejunum and ileum of 3 animals. Both myenteric and submucosal neurons were counted. For embryonic tissue analysis, all cells were counted on a minimum of 3 sections per investigated region of the gut in mutant and littermate WT embryos. Paired Student’s *t* test comparisons were performed for statistical examinations. Counting of fluorescent cells was performed using the “Cell Counter” plugin in ImageJ or directly under a Zeiss fluorescent microscope. In all graphs, bars indicate mean ± Standard Deviation of the Mean (SD).

### Analysis of single cell RNA-sequencing data

#### Clustering analysis

To identify molecularly distinct cell types in juvenile (P21) dataset, we classified the cells into transcriptionally similar clusters. We filtered 9,141 starting cells based on three quality control (QC) covariates: the number of UMI counts per cell, the number of genes per cell and the fraction of counts from mitochondrial genes per cell. Cells with <1,500 genes or >8,000 genes are removed. To mitigate potential doublets, we also excluded cells with >60,000 UMI counts. Of the remaining cells we further removed cells with the fraction of counts from mitochondrial genes <0.1 (for cells with <4000 genes) and <0.25 (for cells with >4000 genes) to account for injured cells. Downstream analysis was performed using the R package *Seurat* version 3.1.2.^8^ Briefly, we normalized the expression data using regularize negative binomial regression method^9^ implemented in the *SCTransform* function while regressing out contribution from mitochondrial gene counts, retaining 3,000 variable genes. Prominent sex specific genes (*Xist, Gm13305, Tsix, Eif253y, Ddx3y, Uty*) and a set of immediate early genes (*Fos, Jun, Junb, Egr1*) were removed from variable genes before principal component (PC) analysis. We then performed preliminary clustering with permissive parameters to identify and remove cluster that are of low quality and are non-enteric. At this level, we constructed K-nearest neighbor graph based on Euclidian distance in 50 principal components (PCs) space and identified distinct communities of cells using Leiden algorithm (res. = 1.5). We removed clusters that shows both high *Plp1* and high *Elavl4* as they likely represented neuron-glia contamination, and clusters that have low UMI counts per cells, low genes per cell and high fraction of mitochondrial genes, as the indication of low quality cells. The remaining cells were subject to second level clustering (35 PCs, res. = 1.0), yielding a clear separation of enteric glia and enteric neuron clusters. Only neuron clusters (4,892 cells in total) were retained for third level clustering (30 PCs, res. = 0.4), yielding 12 final neuron clusters. The clusters were visualized on two dimensional using the *RunUMAP* function (min.dist = 0.5, n_neighbors = 30L, umap.method = “umap-learn”, metric = “correlation”).

For embryonic datasets, we employed a similar approach, but with dataset-specific parameters and Louvain algorithm for clustering. We obtained 3,468 cells from E15.5. After removing cells with <1,000 genes or >6,000, with >40,000 UMI counts (to mitigate potential doublets), and with > 0.05 fraction of mitochondrial genes, we retained 3,260 cells for first level clustering (50 PCs, res. = 0.8). At this step, we removed one non-enteric cluster. The remaining cells were then subject to second level clustering (30 PCs, res. = 0.9), yielding 12 clusters. Similarly, of 2,996 stating cells from E18.5, we removed cells with <1,000 genes or >8,000, with >40,000 UMI counts, and with > 0.1 fraction of mitochondrial genes, yielding 2,733 remaining cells. We removed one non-enteric clusters at first level clustering (50 PCs, res. = 0.8). After second level clustering (30PCs, res. = 0.9), we obtained 14 clusters. We then performed differential expression analysis and, based on differential expressed genes, categorized each cluster as one of the following major cell types: neuron (*Elavl4, Tubb3*), neuroblast (*Ascl1*), progenitor (*Sox10*), glia (high *Plp1*), and schwann cell precursor cells (*Dhh*). Clusters with similar type were merged before re-analyzing differential expression to identify major gene modules (Figure 5). The clusters were visualized on two dimensional using the *RunUMAP* function (for E15.5, min.dist = 0.48, n_neighbors = 69L, umap.method = “umap-learn”, metric = “correlation”; for E18.5, min.dist = 0.5, n_neighbors = 49L, umap.method = “umap-learn”, metric = “correlation”).

#### Differential expression analysis

To identify cluster-enriched genes, we performed non-parametric Wilcoxon rank sum test for cells in each cluster compared to all other cells. p-values were adjusted based on Bonferroni correction, giving false discovery rate (FDR). Genes with FDR < 0.01, at least 0.25 average fold change (log-scale) and at least 25% cluster-specific detection (percent of cells expressing a particular gene in a cluster) were defined as enriched genes for each cluster. Top genes were then scrutinized in order of average fold change (log-scale).

#### Supervised cell type assignment

We used the hierarchical statistical framework *CellAssign*^10^ to compute the probability that each cell belongs to a cell type defined by a set of know makers genes. We used priori makers information gathered from previous studies. The curated markers were converted to binary codes before being used as an input to the model. Cell specific size factors were computed from the *sizeFactor* function of the R package *scarn*. Individual cells were annotated to the cell type with maximum probability. For Figure 1d and 1e, the following priori markers were used; *Elavl4, Tubb3, Stmn2, Snap25* (enteric neuron); *Sox10, Plp1, S100b* (enteric glia); *Ube2c, Top2a, Cenpef* (cycling); *Actg2, Mylk* (smooth muscle). For presentation, we combined cells assigned to “cycling”, “smooth muscle” and “unassigned” to a new group, “Other”. Also note that, for comparison of cell types obtained from *Wnt1Cre;R26Tomato* and *Baf53bCre;R26Tomato* experiments (Fig. 1e), we applied the same permissive pre-processing thresholding (UMI counts > 600) for both datasets. For Figure 1i and 1j, priori markers were curated based on a series of immunohistochemical studies on the mouse small intestine.^11–13^ The following markers were used: *Calca, Calcb, Chat, Slc18a3, Nefm, Calb1, Calb2* (IPAN); *Chat, Slc18a3, Calb2, Tac1* (excitatory motor neuron); *Nos1, Vip, Npy* (inhibitory motor neuron); *Chat, Slc18a3, Nos1, Vip* (interneuron 1); *Chat, Slc8a3, Calb2, Sst* (interneuron 2); *Chat, Slc18a3, Ddc, Slc6a4* (interneuron 3).

#### Cell cycle assignment

For each cell, we computed a score based on its expression of G2/M and S phase markers; cells that express neither of these markers are likely not cycling or are in G0/G1 phase. The scoring strategy is described in Tirosh et al. 2016.^14^ We used *Seurat*’s implementation of this strategy in the *CellCycleScoring()* function.

#### Label transfer

For comparison of our newly proposed enteric neuron clusters to our previously reported ones (Extended Data Fig. 1b), we used expression data and cluster identities of enteric neurons at level L6 which can be found at http://mousebrain.org. Accession numbers for raw sequence data from *Wnt1Cre;R26RTomato* P21 animals is SRP135960, available at https://www.ncbi.nlm.nih.gov/sra/SRP135960). Both datasets (*Wnt1Cre;R26RTomato* and *Baf53bCre;R26RTomato*) were subjected to the same pre-processing steps using regularize negative binomial regression method^9^, while retaining original cluster identities, before undergoing *Seurat*’s label transfer^15^, using the *FindTransferAnchors()* function with 30 PCs and then the *TransferData()* function with *Baf53bCre;R26Tomato* dataset as the reference dataset. Finally, the mapping identities were presented as Alluvial plot using R package *ggallivial*.

For mapping of juvenile cell types to embryonic datasets (Fig. 6a and 6b), we employed similar approach as mentioned above. We compiled a juvenile reference dataset by combining enteric neurons from third level clustering (retaining their ENC1-12 identities) with enteric glial cells from second level clustering (removing cells with high expression of *Elavl4* and *Tubb3*, and merging all subclass to ‘enteric glia’), giving a total of 13 class identities (12 ENCs and enteric glia). Here, we used 30 PCs for mapping of E15.5 dataset and 40 PCs for mapping of E18.5 dataset. Only cells with >0.5 maximum prediction score were mapped.

#### Computational screening of HGNC gene sets

To assess whether given gene sets can explain heterogeneity within the 12 ENCs, we performed a computational screening as previously described^16^, which is based on the *MetaNeighbor* method.^17^ In brief, we defined a Spearmann correlation subnetwork of cells in our dataset by their expression of a given gene set, and performed a stratified cross-validation. In doing so, we divided our dataset into test and train sets and then held back cluster identity from one set at a time and predicted based on the other. The efficacy of cells of the same cluster were recovered for each cluster and reported as the area under the receiver operator characteristic curve (AUROC). We then ranked the gene sets by the mean AUROC. We perform this test on HGNC^18^ (HUGO Gene Nomenclature Committee) families curated by Paul et al. 2017^16^, selecting only gene sets with more than three detectable genes.

#### Partition-based graph abstraction and pseudotime analysis

We used partitioned-based graph abstraction (PAGA) for interring developmental trajectories.^19^ We found this approach most suitable with our developmental datasets which consist of both cycling and branching topology. We used cluster identities from Louvain algorithm (as mentioned above) to define partitions of the graph of neighborhood relation among data points. PAGA was computed using the *tl.paga()* function implemented in the Python package *Scanpy 1.4.5.1*^20^. For pseudotime inference, we employed diffusion pseudotime method which infers progression of cells through geodesic distance along the graph.^21^ We used *Scanpy*’s *tl.dpt(),* setting the root at the cluster that most likely correspond to neuroblast, and the number of diffusion components to 10. Together with PAGA, which allowed detection of Branches, we were able to define developmental paths within each of developmental dataset. Cells were then re-ordered within each path before plotting gene expression (Fig. 6g).

