## Extended Data Figure 1 for "Diversification of molecularly defined myenteric neuron classes revealed by single cell RNA-sequencing"

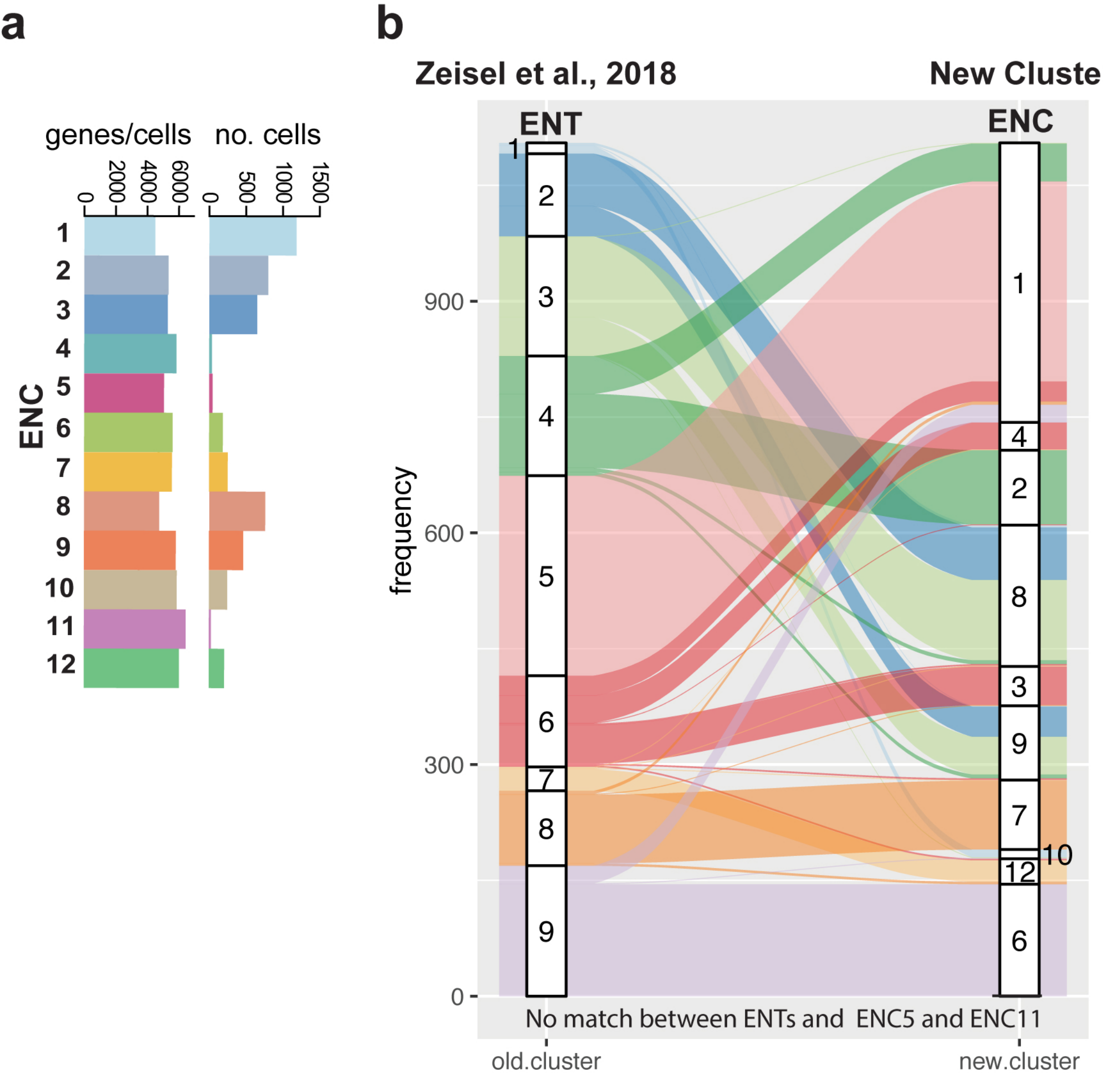


**Extended Data Figure 1: Supportive Data related to Figure 1e-f.**
a) Table showing number of cells and average number of genes per cell in each ENC. b) Label transfer plot indicating relation between cluster ENT1-9 found in Zeisel et al., 2018 and cluster ENC1-12 presented in this study. Notably, ENC5 (*Sst*) and ENC11 (*Npy/Th/Dbh*) represent new clusters not present in the Zeisel et al., 2018 data set. ENTs representing plausible exitatory (ENT4-6) and inhibitory (ENT2,3) motor neurons were not retained, but clustered differentially into ENC1-4 and ENC8-9, respectively. Nomenclature: ENT: Enteric Neuron Type; ENC: Enteric Neuron Class
