## Extended Data Figure 2 for "Diversification of molecularly defined myenteric neuron classes revealed by single cell RNA-sequencing"

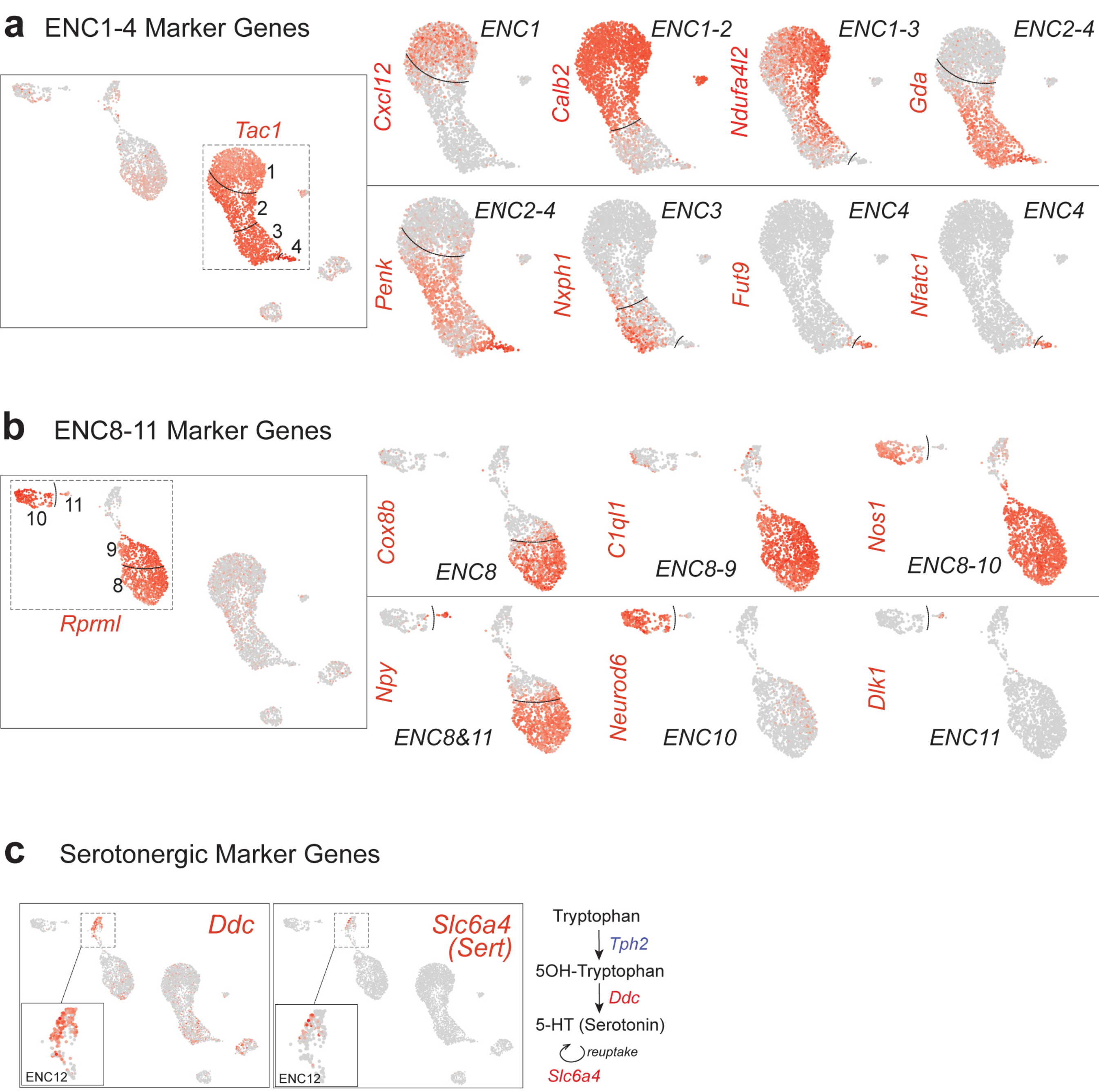


**Extended Data Figure 2: Gene expression related to Figure 1h.**
Feature plots displaying common and diverge gene expression in:
a) *Tac1*$^+$ clusters (ENC1-4)
b) *Rprml*$^+$ clusters (ENC8-11).
c) Feature plots displaying expression of genes correlated to serotonin production (*Ddc*) and re-uptake (*Slc6a4*) in ENC12.
