## Extended Data Figure 3 for "Diversification of molecularly defined myenteric neuron classes revealed by single cell RNA-sequencing"

#### a) Neurotransmission

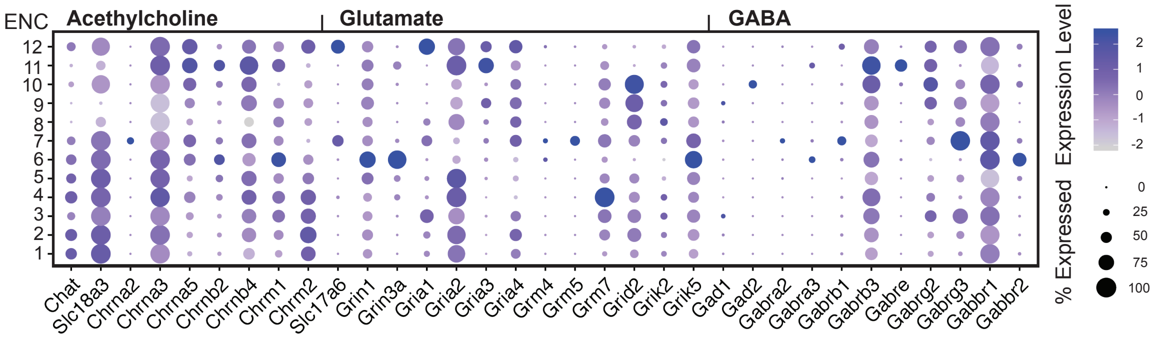


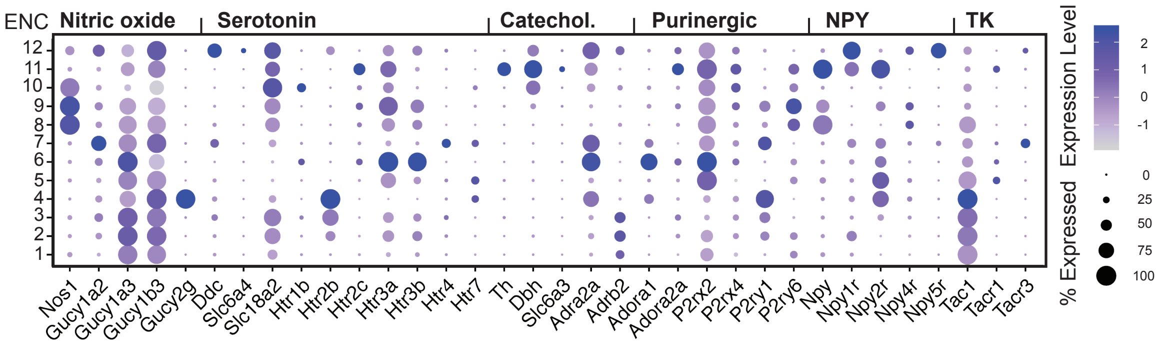


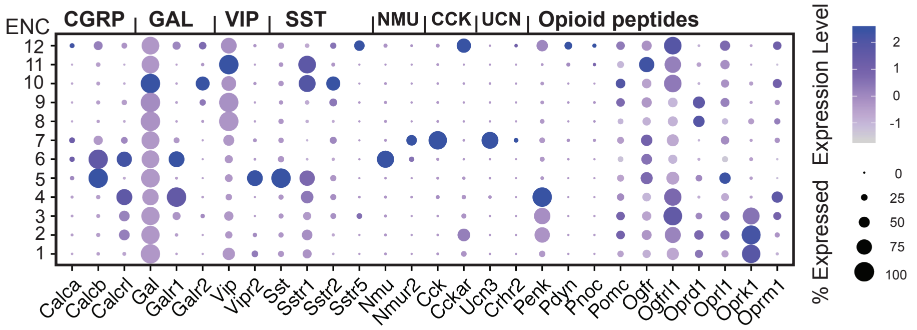

### b) Cell-Cell Signaling

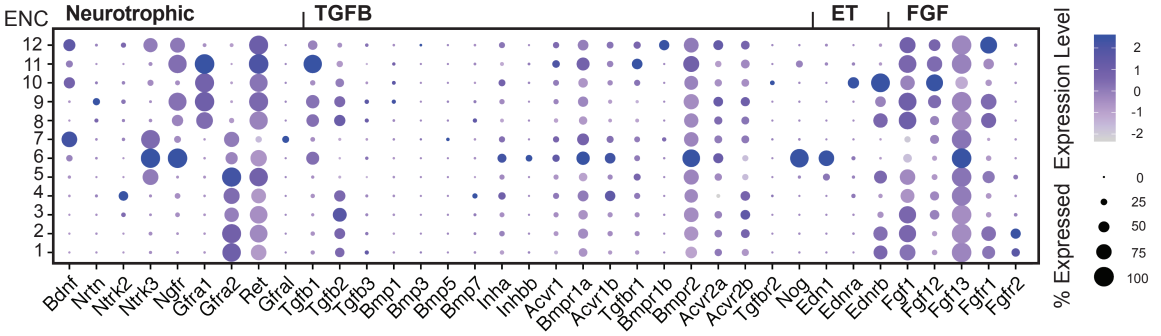


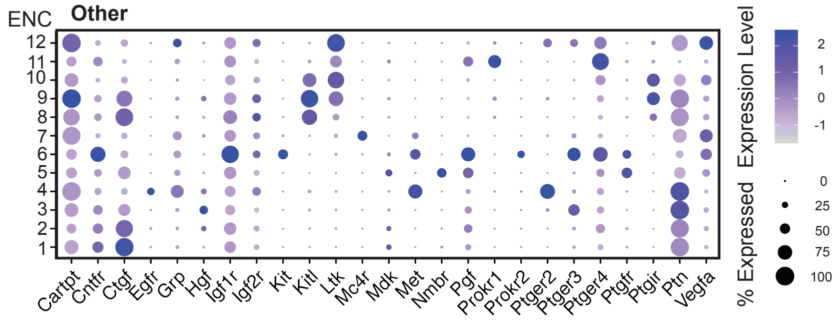

#### c) Transcription Factors

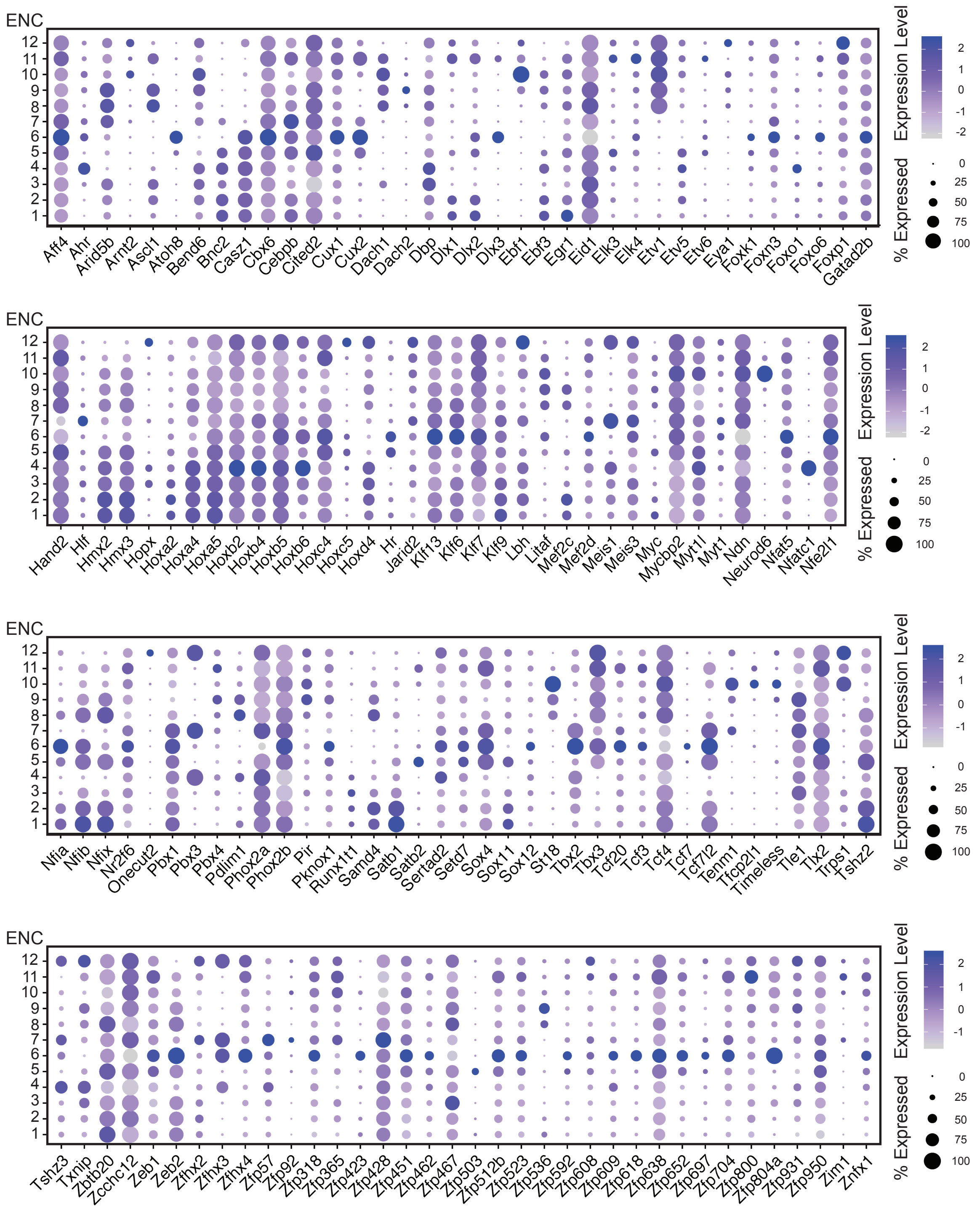

#### d) Adhesion Molecules

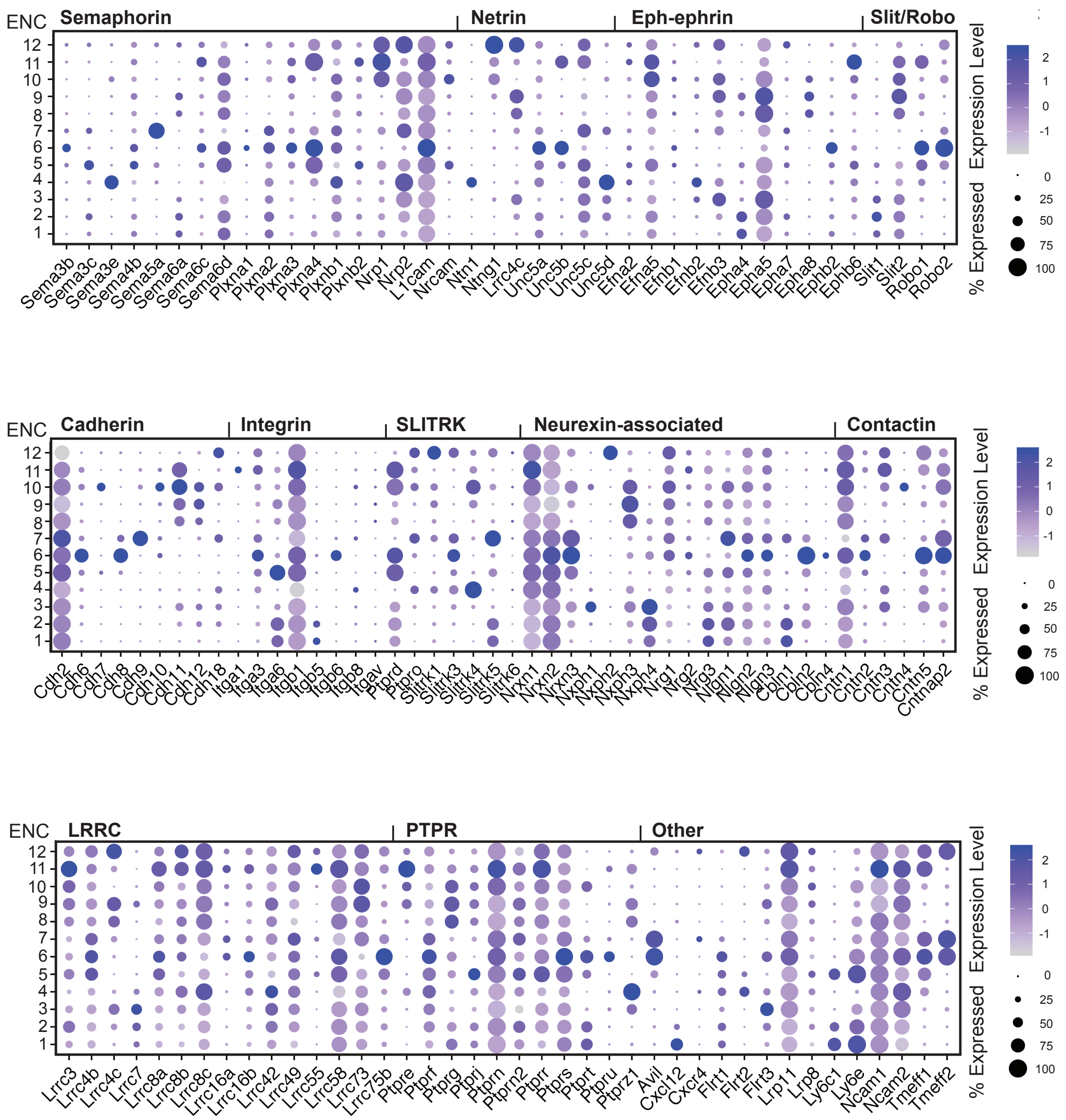

### e) Ion Channels

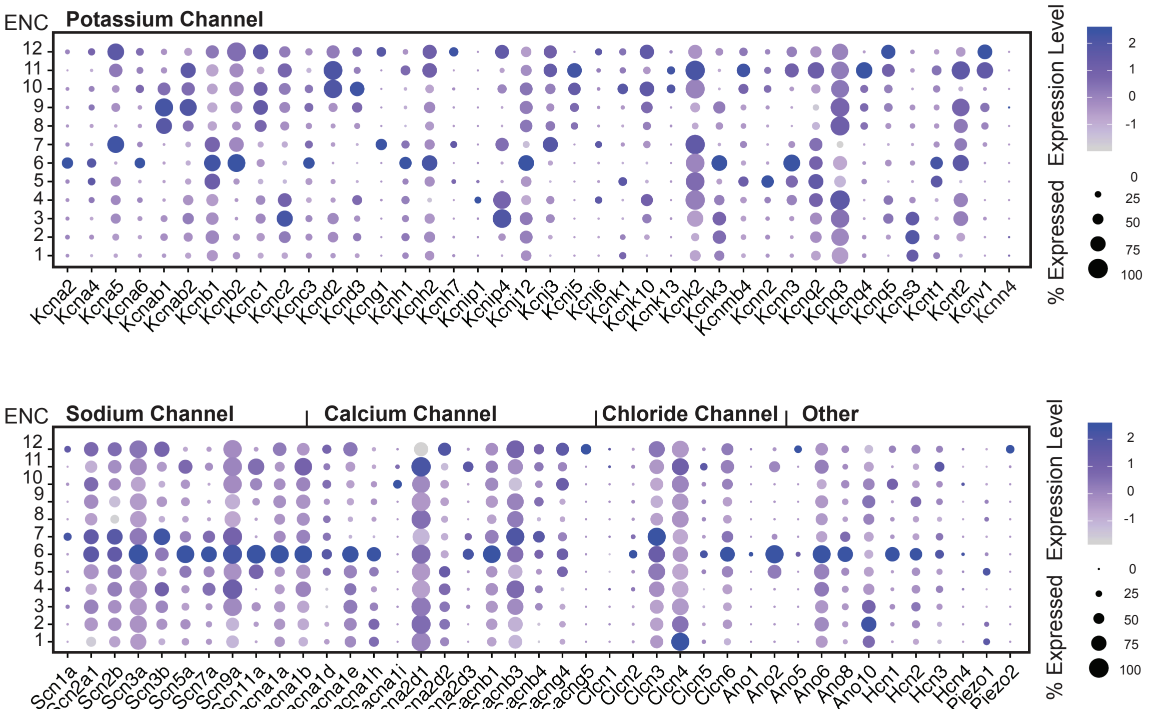


### f) Membrane Trafficking

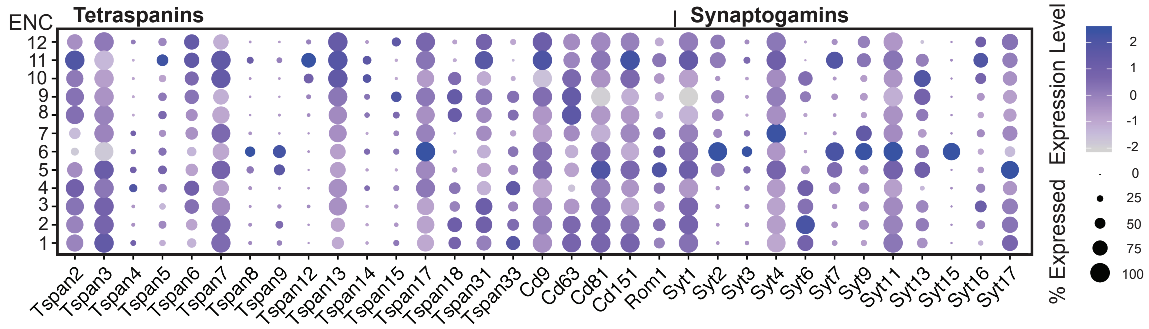


**Extended Data Figure 3. Dot plots displaying expression of genes conferring neuronal phenotypes in ENC1-12.**
Gene categories involved in a) neurotransmission, b) other cell-cell signaling, c) transcription, d) adhesion e) ion transport, and f) membrane trafficking.
Color scale represents z-score and dot size represent percent of cells with non-zero expression within a given class. ENC: Enteric Neuron Class; GABA: Gamma-aminobutyric acid; NPY: Neuropeptide Y; TK:Tachykinin; CGRP: Calcitonin gene-related peptide; GAL:Galanin;VIP: Vasoactive intestinal peptide; SST: Somatostatin; NMU: Neuromedin U; CCK: Cholecystokinin; UCN: Urocortin; TGFB: Transforming growth factor beta; ET:Endothelin; FGF: Fibroblast growth factor LRRC: Leucine Rich Repeat Containing; PTPR: Protein tyrosine phosphatase receptor.
