## Extended Data Figure 4 for "Diversification of molecularly defined myenteric neuron classes revealed by single cell RNA-sequencing"

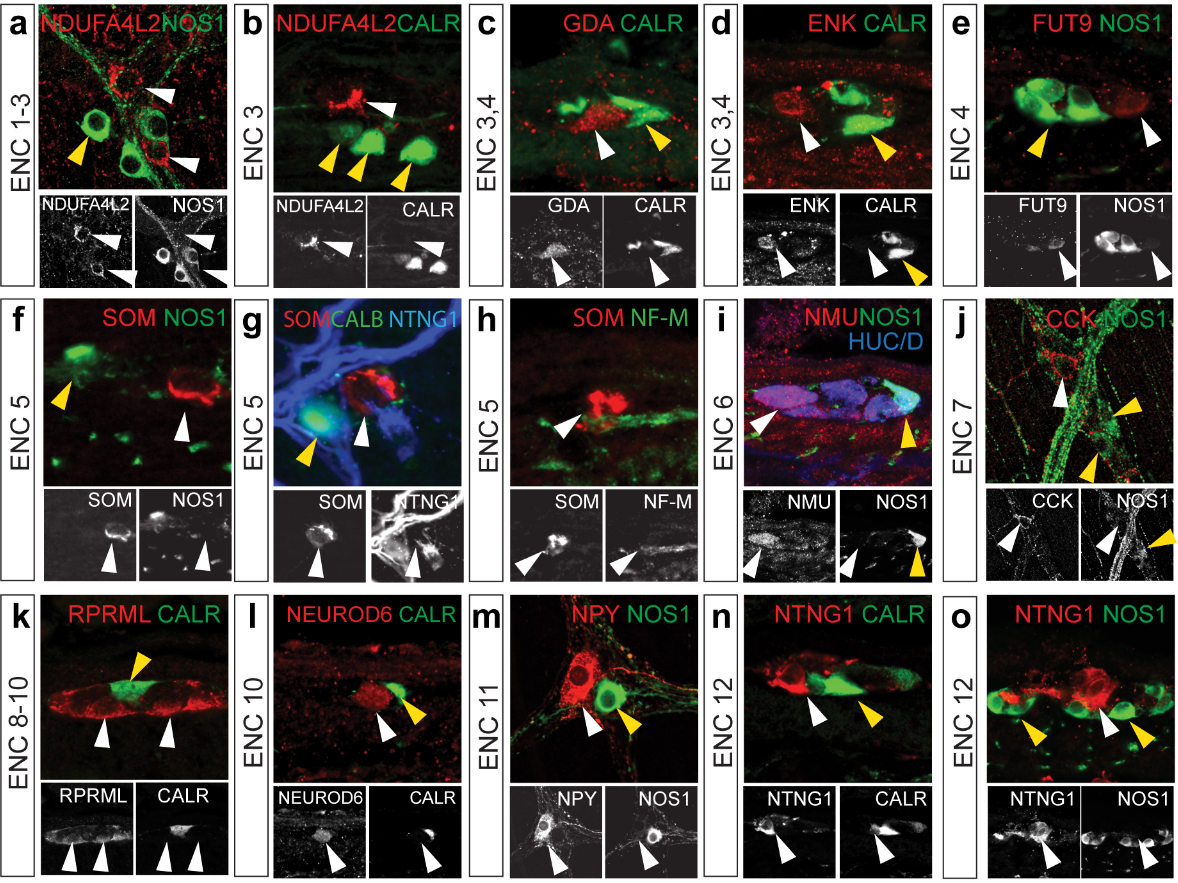


**p** ENC markers confirmed by immunohistochemistry

| | POSITIVE | NEGATIVE |
|---|---|---|
| ENC 1 | NDUFA4L2 CALR SP | NOS1 |
| ENC 2 | NDUFA4L2 CALR ENK GDA SP | NOS1 |
| ENC 3 | NDUFA4L2 ENK GDA SP | NOS1 CALR |
| ENC 4 | ENK GDA **FUT9** SP | NOS1 CALR |
| ENC 5 | **SOM** CALR CGRP | NTNG1, CALB NOS1, NF-M |
| ENC 6 | **NMU** CGRP NF-M$^{+/-}$ NOGGIN CALR (CALB) | NOS1 |
| ENC 7 | **CCK UCN3** NF-M$^{+/-}$ CALB$^{FEW}$ VGLUT2 (CGRP) | NOS1 5-HT |
| ENC 8 | NOS1 GAL RPRML NPY | CALR |
| ENC 9 | NOS1 GAL RPRML | NPY |
| ENC 10 | **NEUROD6 GAD2** NOS1 RPRML NF-M, GAL | CALR |
| ENC 11 | NPY$^{high}$ CALR TH DBH | NOS1 |
| ENC 12 | **NXPH2** NTNG1 CALB NF-M ENK$^{+/-}$ **PIEZO2**$^{+/-}$ **5-HT**$^{+/-}$ VGLUT2 | CALR, CGRP NOS1 |

**BOLD**: Unique Marker

**Extended Data Figure 4: Confirmation of negative markers for ENC1-12 and summary table of all validated ENC markers.** a-o) Immunohistochemical localisation of positive and negative marker proteins. Pictures show either myenteric peel preparations or transverse sections at P21-P90. p) Table summarizing ENC markers verified by immunohistochemistry. white arrowhead: positive marker;yellow arrowhead: negative marker. CALB: Calbindin; CALR: Calretinin; CCK: Cholecystokinin; CGRP:Calcitonin Gene Related Peptide DBH:Dopamine Beta Hydroxylase; ENK: Enkephalin; GAD2: Glutamic Acid Decarboxylase 2;GAL: Galanin; NF-M: Neurofilament M; NMU: Neuromedin U; NOS1; Nitric Oxide Synthase 1; NPY: Neuropeptide Y; NTNG1: Netrin G1; RPRML: Reprimo-Like; SOM: Somatostatin; SP: Substance P; TH: Tyrosine Hydroxylase; VGLUT2: Vesicular Glutamate Transporter 2
