## Extended Data Figure 5 for "Diversification of molecularly defined myenteric neuron classes revealed by single cell RNA-sequencing"

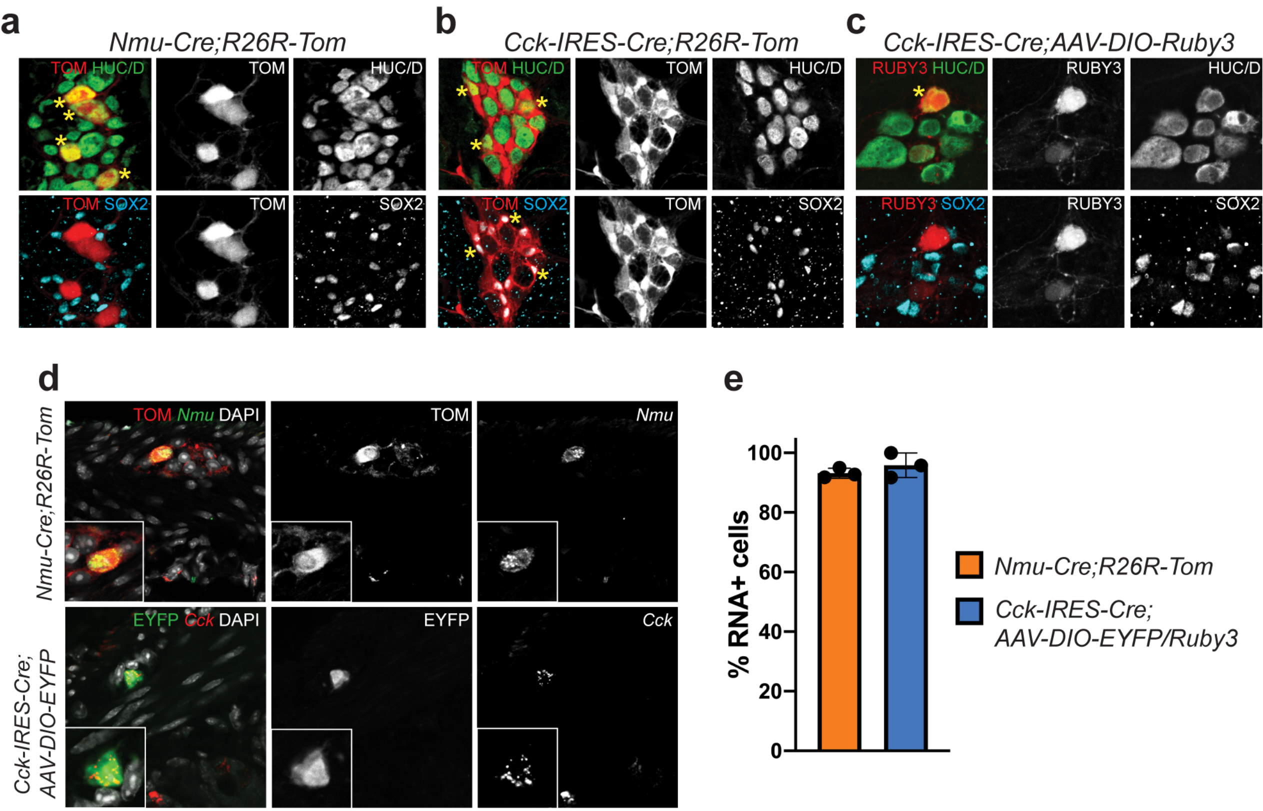


**Extended Data Figure 5: Validation of transgenic Cre-lines for the investigation of ENC6 and ENC7.** a) Myenteric plexus peel from *Nmu-Cre;R26R-Tom* mouse showing TOM in HUC/D$^+$ neurons (stars) and its exclusion from enteric glia (SOX2$^+$). b) Myenteric plexus peel from *Cck-IRES-Cre;R26-Tom* mouse showing TOM in both neurons (stars) and enteric glia (stars). c) Myenteric plexus peel of *Cck-IRES-Cre* mouse injected with *AAV-DIO-Ruby3* showing RUBY3 only in neurons and not in glia. d) Transverse sections showing that *Nmu* and *Cck* RNA expression correlate with reporter$^+$ neurons in *Nmu-Cre;R26R-Tom* and *Cck-IRES-Cre; AAV-DIO-EYFP/Ruby3* animals. e) Graph showing the percentage of reporter$^+$ neurons expressing the reciprocal RNA. A total of 638 reporter$^+$ neurons were investigated in three *Nmu-Cre;R26R-Tom* mice, and 71 reporter$^+$ neurons were investigated in three *Cck-IRES-Cre;AAV-DIO-EYFP/Ruby3* mice.TOM: dtTomato
