## Extended Data Figure 6 for "Diversification of molecularly defined myenteric neuron classes revealed by single cell RNA-sequencing"

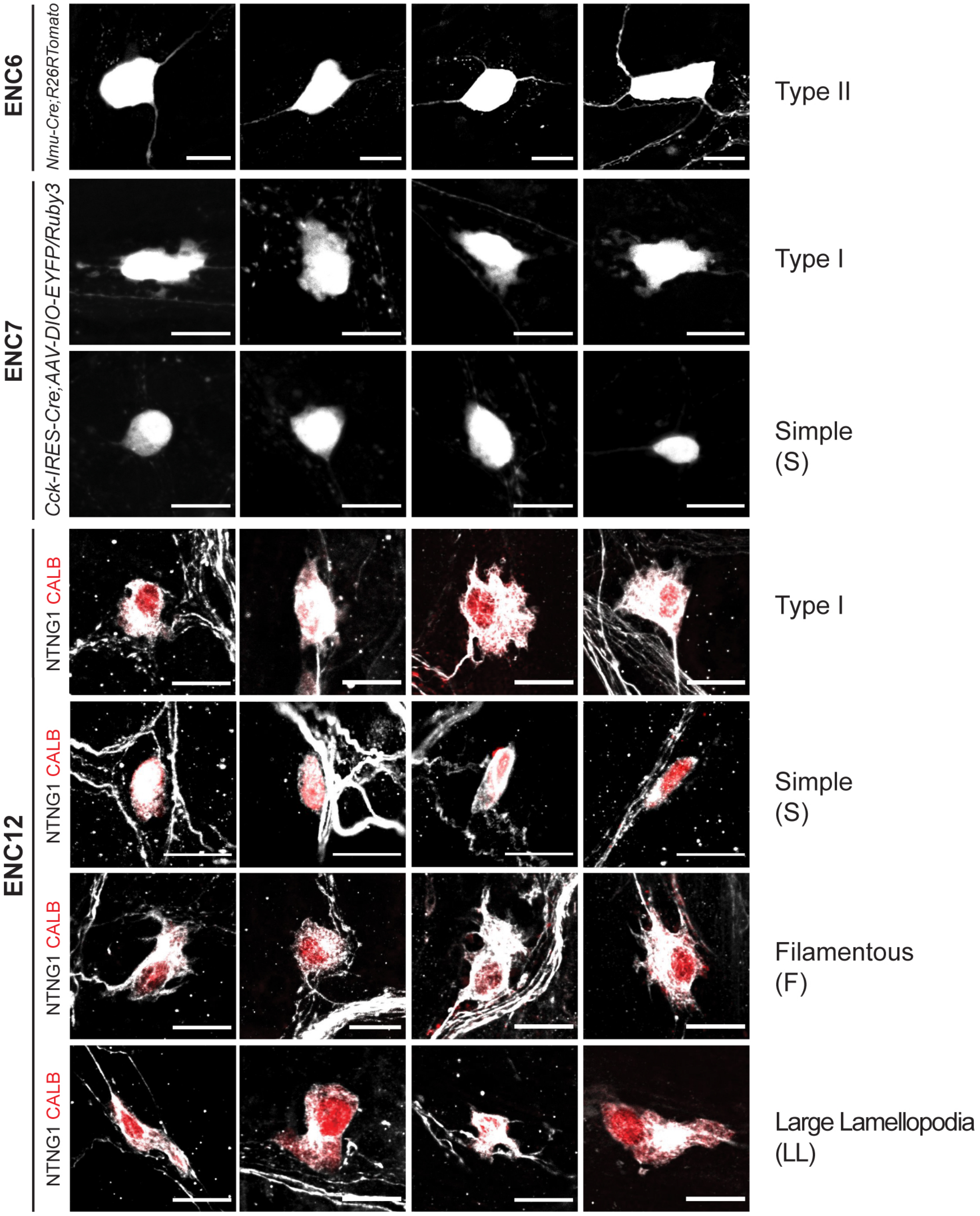


**Extended Data Figure 6: Morphological characterization of ENC6, 7 and 12.** Related to Figure 4e-g. Representative examples of each morphological type found within ENC6, 7 and 12. Scale bar indicates 20$\mu$m.
