## Extended Data Figure 7 for "Diversification of molecularly defined myenteric neuron classes revealed by single cell RNA-sequencing"

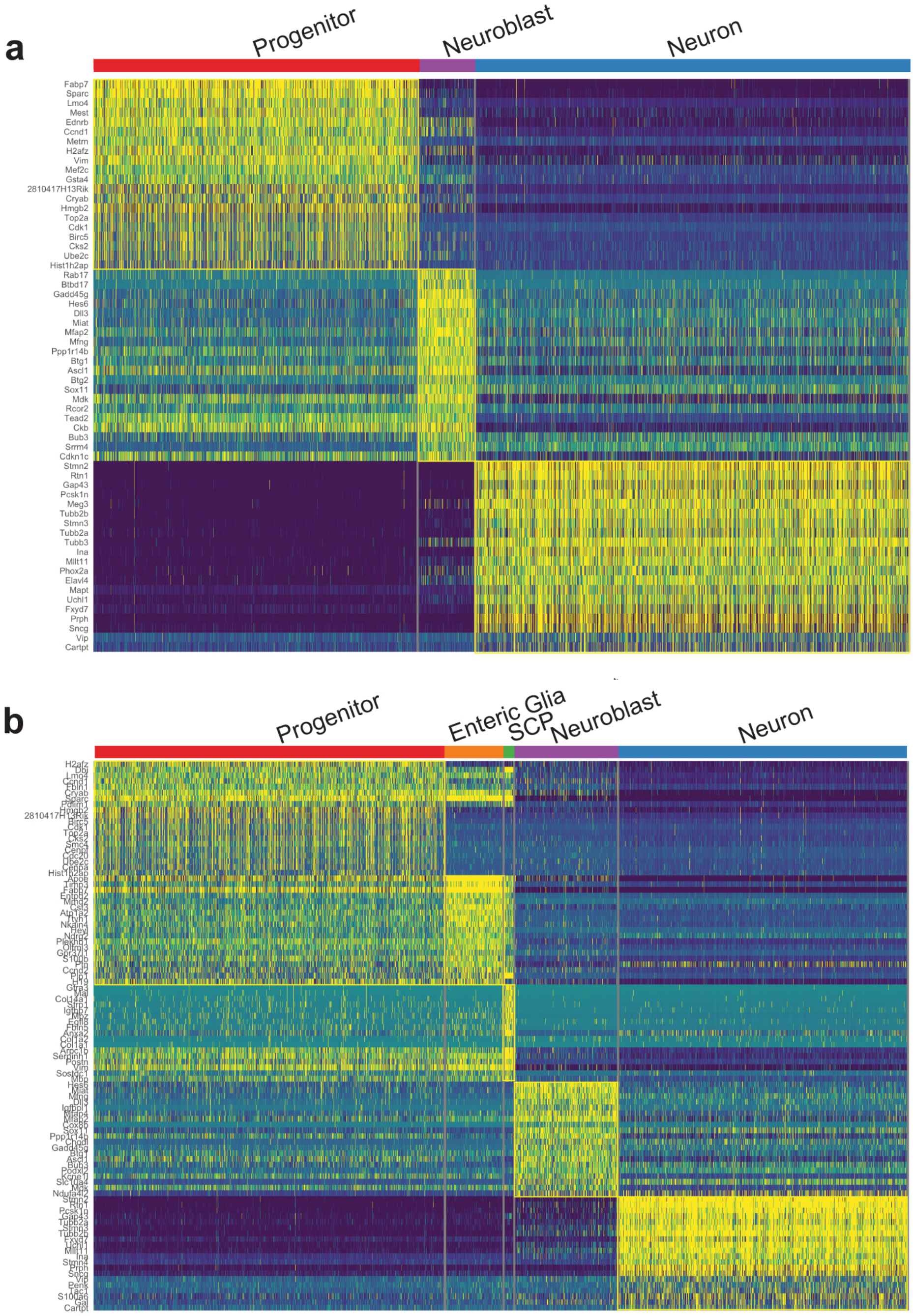


**Extended Data Figure 7: Heatmaps showing top marker genes for clusters representing generic cell states of the developing ENS.** Related to Figure 5. a) E15.5 b) E18.5. SCP: Schwann Cell Precursor
