## Extended Data Figure 8 for "Diversification of molecularly defined myenteric neuron classes revealed by single cell RNA-sequencing"

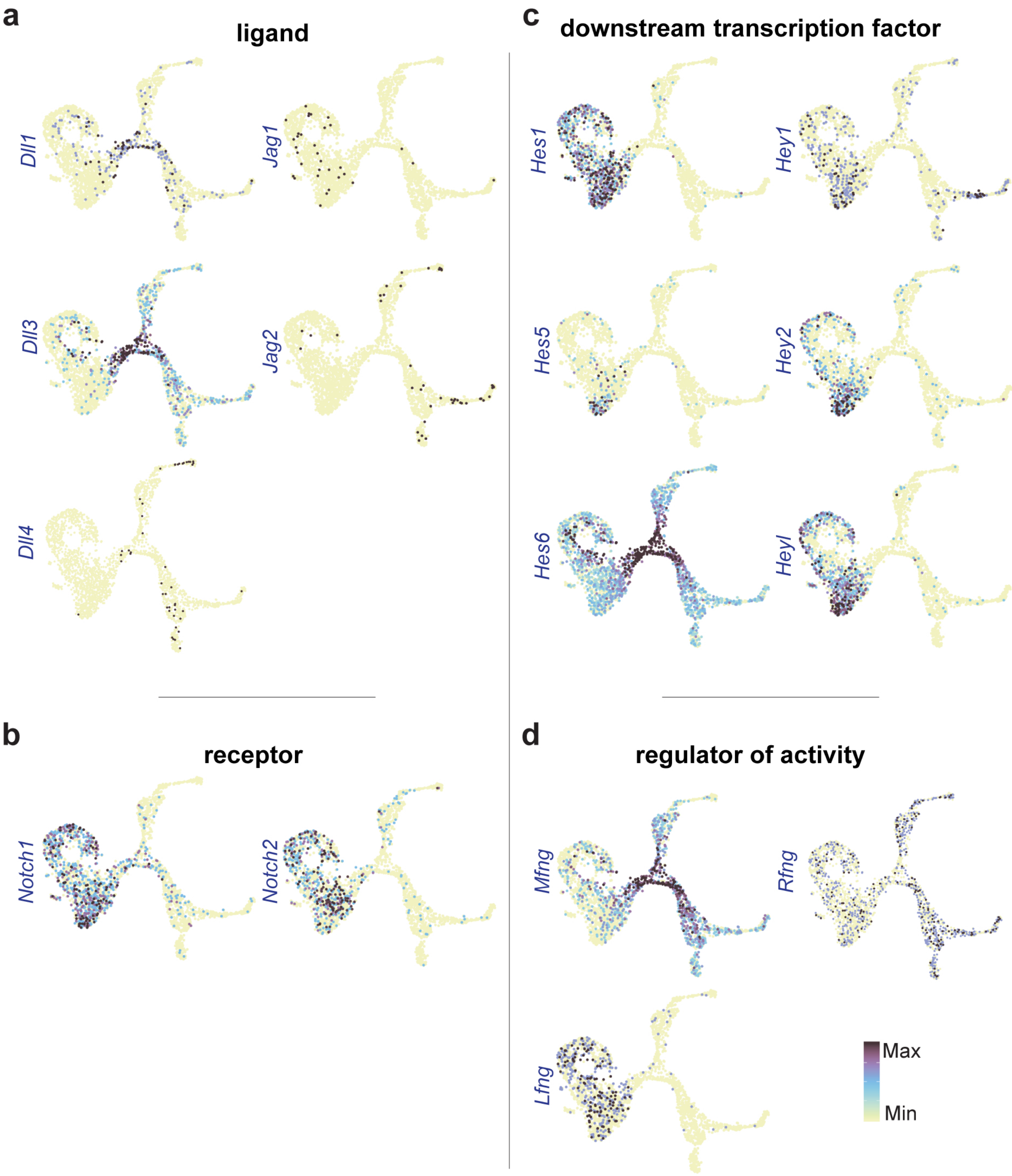


**Extended Data Figure 8: Feature plots of ENS at E18.5 displaying expression of Notch signaling genes. Related to Figure 5**

a) Ligands; note the predominant expression of *Dll1* and *Dll3* in neuroblasts

b) Receptors; note the predominant expression of Notch1,2 in progenitors.

c) Downstream transcription factor; note the enriched expression of *Hes6* in neuroblasts, *Hes1* in progenitors and *Hes5*, *Heyl* and *Hey2* in enteric glia.

d) Regulator of activity; note the enriched expression of *Mfng* in neuroblasts and Lfng in progenitors.
