## Supplementary Table 2 for "Diversification of molecularly defined myenteric neuron classes revealed by single cell RNA-sequencing"

#### Primary Antibodies used in the study

| Target | Host | Dilution | Source | Comment: |
|---|---|---|---|---|
| 5-HT | Goat | 1:500 | Abcam ab66047 | |
| ANO1 | Rabbit | 1:1,000 | Abcam ab64085 | |
| ANO2 | Mouse | 1:1,000 | Santa Cruz sc-390956 | |
| CALB | Rabbit | 1:1,000 | Chemicon AB1778 | recognises Calretinin |
| CALB | Rabbit | 1:500 | Swant CB38a | recognises Calretinin |
| CALB | Mouse | 1:500 | Swant CB300 | |
| CALB | Goat | 1:500 | R&D AF3320 | recognises Calretinin |
| CALR | Mouse | 1:500 | Santa Cruz sc-365956 | |
| CALR | Rabbit | 1:1,000 | Chemicon AB5054 | |
| CCK | Rabbit | 1:300 | Abcam ab83180 | enhanced with colchicine |
| CCK | Rabbit | 1:16,000 | Immunostar 20078 | enhanced with colchicine |
| CGRP | Goat | 1:1,500 | AdBsero 1720-9007 | enhanced with colchicine |
| CGRP | Rabbit | 1:5,000 | Sigma C8198 | enhanced with colchicine |
| DBH | Rabbit | 1:1,000 | Immunostar 22806 | |
| ENK | Rabbit | 1:3-6,000 | St John's STJ98660 | |
| ENK | Mouse | 1:300 | Abcam ab150346 | |
| FUT9 | Rabbit | 1:300 | Novus Bio NBP2-57960 | |
| GAD2 | Rabbit | 1:50,000 | Kind gift from J.Kaltschmidt | |
| GAL | Goat | 1:200 | Abcam ab99452 | Weak staining in cell bodies |
| GDA | Rabbit | 1:1000 | Atlas Antibodies HPA024099 | |
| GFP | Goat | 1:1,000 | Abcam ab6662 | FITC-conjugated |
| HuC/D | Mouse | 1:300 | Molecular Probes A21271 | |
| NDUFA4L2 | Rabbit | 1:200 | St John's STJ94374 | Weak staining in cell bodies |
| NDUFA4L2 | Rabbit | 1:5,000 | St John's STJ116500 | Weak staining in cell bodies |
| NEUROD6 | Rabbit | 1:100 | Abcam ab85824 | |
| NF-M | Mouse | 1:500 | Abcam ab7794 | |
| NMU | Mouse | 1:200 | Santa Cruz sc-398600 | Variable weak |
| NMU | Rabbit | 1:100 | St John's STJ94433 | unspecific |
| NMU | Rabbit | 1:200 | St John's STJ116312 | Variable weak, only on sections |
| NOS1 | Goat | 1:1,000 | Abcam ab1376 | |
| NOS1 | Rabbit | 1:200 | Santa Cruz sc-648 | |
| NOS1 | Mouse | 1:300 | Santa Cruz sc-5302 | |
| NPY | Rabbit | 1:3,000 | Immunostar 22940 | |
| NTNG1 | Rabbit | 1:200 | Abcam ab221456 | |
| NTNG1 | Mouse | 1:400 | Santa Cruz sc-271774 | |
| NXPH2 | Rabbit | 1:200 | Atlas Antibodies HPA034759 | |
| PBX1 | Rabbit | 1:1000 | Cell Signaling #4542 | |
| PBX3 | Rabbit | 1:200 | Sigma AV32070 | |
| PGP9.5 | Rabbit | 1:1,000 | ThermoFisher PA5-29012 | |
| PGP9.5 | Mouse | 1:300 | Novus Bio NB600-1160 | |
| PIEZO2 | Rabbit | 1:200 | LifeSpan LS-C180178 | mainly in processes |
| Pro-CCK | Rabbit | 1:800 | Frontiers Institute AB2571674 | |
| RFP/Tomato /Ruby | Rat | 1:1,000 | Chromotek, 5F8 | |
| RPRML | Rabbit | 1:3,000 | Atlas Antibodies HPA062668 | |

| | | | | |
|---|---|---|---|---|
| SOM | Rat | 1:300 | Merck MAB354 | |
| Sox10 | Goat | 1:2,000 | R&D Systems AF2864 | |
| Sox2 | Rabbit | 1:4,000 | Seven Hills WRAB-1236 | tissue becomes dotty |
| Substance P | Rabbit | 1:1,000 | Chemicon 1566 | |
| TH | Rabbit | 1:500 | Pelfreeze P40101-0 | |
| TH | Sheep | 1:300 | Novus Bio NB300-110 | |
| UCN3 | Mouse | 1:200 | Santa Cruz sc-517449 | Unspecific? |
| UCN3 | Rabbit | 1:1,000 | Biometrik CAU23545 | |
| UCN3 | Goat | 1:200 | St Johns STJ71448 | |
| VGLUT2 | Guinea pig | 1:5,000 | Millipore AB2251 | Enhanced by colchicine |
| VIP | Rabbit | 1:200 | AbDSero 95350204 | |

Good staining
OK staining
Weak staining
Unspecific/undetected staining

### Secondary antibodies used in the study

| **Alexa Fluor 488/FITC (dilution 1:400)** | **Alexa Fluor 555/594 (dilution 1:1,000)** | **Alexa Fluor 647/Cy5 (dilution 1:250)** |
|---|---|---|
| Donkey anti-goat 488<br>ThermoFisher A11055 | Donkey anti-goat 555<br>ThermoFisher A21432 | Donkey anti-goat 647<br>ThermoFisher A21447 |
| Donkey anti-guinea pig 488<br>Jackson 706-545-148 | Goat anti-guinea pig 555<br>ThermoFisher A21435 | Donkey anti-guinea pig Cy5<br>Jackson 706-175-148 |
| Donkey anti-mouse 488<br>ThermoFisher A21202 | Donkey anti-mouse 555<br>ThermoFisher A31570 | Donkey anti-mouse 647<br>ThermoFisher A31571 |
| Goat anti-mouse IgG1 488<br>ThermoFisher A21121 | Goat anti-mouse IgG1 555<br>ThermoFisher A21127 | Goat anti-mouse IgG1 647<br>ThermoFisher A21240 |
| Goat anti-mouse IgG2a 488<br>ThermoFisher A21131 | Goat anti-mouse IgG2a 555<br>ThermoFisher A21137 | Goat anti-mouse IgG2a 647<br>ThermoFisher A21241 |
| Goat anti-mouse IgG2b 488<br>ThermoFisher A21141 | | Goat anti-mouse IgG2b 647<br>ThermoFisher A21242 |
| Rat anti-mouse IgG1 FITC<br>ThermoFisher 11-4015-82 | | |
| Donkey anti-rat 488<br>ThermoFisher A21209 | Donkey anti-rat 594<br>ThermoFisher A21208 | Goat anti-rat 647<br>Cell Signaling 4418S |
| Donkey anti-rabbit 488<br>ThermoFisher A32790 | Donkey anti-rabbit 555<br>ThermoFisher A31572 | Donkey anti-rabbit 647<br>ThermoFisher A31573 |
| Donkey anti-sheep 488<br>ThermoFisher A11015 | Donkey anti-sheep 555<br>ThermoFisher A21436 | |
